# Amyloid precursor protein causes fusion of promyelocytic leukemia nuclear bodies in human hippocampal areas with high plaque load

**DOI:** 10.1101/2020.04.16.001255

**Authors:** David Marks, Natalie Heinen, Lisa Bachmann, Sophia Meermeyer, Michelle Werner, Lucia Gallego, Stephanie Nolte, Peter Hemmerich, Verian Bader, Konstanze Winklhofer, Elisabeth Schröder, Shirley K. Knauer, Thorsten Müller

**Author notes:** shared first authors (David Marks, Natalie Heinen, Lisa Bachmann).

## Abstract

The amyloid precursor protein (APP) is a type I transmembrane protein with unknown physiological function but potential impact in neurodegeneration. The current study demonstrates that APP signals to the nucleus causing the generation of aggregates comprising its adapter protein FE65 and the tumour suppressor proteins p53 and PML. The PML nuclear body generation, known to be of relevance in virus defence and cell division, is induced and fusion occurs over time depending on APP signalling. We further show that the nuclear aggregates of APP C-terminal (APP-CT) fragments together with PML and FE65 are present in the aged human brain but not in cerebral organoids differentiated from iPS cells. Notably, human Alzheimer’s disease brains reveal a highly significant loss of these nuclear aggregates in areas with high plaque load compared to plaque-free areas of the same individual. Based on these results we conclude that APP-CT signalling to the nucleus takes place in the aged human brain and is potentially involved in the pathophysiology of AD. Taken the current knowledge on PML bodies into account, we hypothesize a new role for APP as a twofold virus response protein. The APP-dependent defence strategy includes Aß-virus interaction at the extracellular matrix and APP-CT driven PML aggregation in the nucleus to encapsulate the viral nucleic acid. This defence strategy preferentially occurs in high-plaque regions of the human brain and overstimulation of this pathway results in a pyrrhic victory.

## Introduction

Increased amyloidogenic processing of the amyloid precursor protein (APP) occurs in sporadic Alzheimer’s disease (AD) ^1^, in familial AD with mutations in APP or in its processing enzymes ^2^, and in trisomy 21 patients ^3^. Affected patients suffer from progressive cognitive decline and memory loss, pointing to the fact that APP cleavage is a central mechanism in the pathophysiology of these disorders. Amyloidogenic APP processing causes the generation of three fragments: i) the secreted extracellular domain (sAPPβ), ii) the ß-amyloid peptide (Aî), and iii) the APP C-terminal fragment (APP-CT). The secreted fragment (i) was reported to provoke neurotrophic effects ^4^, Aß (ii) is the main component of amyloidogenic plaques and research over the last decades resulted in the formulation of the amyloid cascade hypothesis ^5^. Finally, APP-CT (iii) was recently suggested to play an important role in a nuclear signal transduction pathway ^6^. The amyloid cascade hypothesis has led to a number of therapeutical approaches, all of which however failed so far in clinical trials ^7^. In contrast, the other two APP cleavage fragments have been less extensively studied. APP-CT indeed is a remarkable protein fragment as it is intrinsically unstructured ^8^. Though, this changes upon interaction with FE65, causing APP-CT to fold into a three-dimensional conformation that can be analysed by x-ray crystallography. Besides APP-CT’s ability to bind a large number of other proteins at the membrane, it was shown that APP-CT is capable to enter the nucleus to subsequently establish a protein complex consisting of additional proteins like FE65, TIP60, and BLM ^9^. The complex is highly dynamic and, over time, the single aggregates fuse together to form large structures of unknown function. The presence of the histone acetyl transferase (TIP60) and the DNA helicase (BLM) in the complex points to a functional role in essential biological mechanisms such as gene expression, DNA replication/damage/repair or chromatin modification. Indeed, a variety of target genes like GSK3β, IDE, and APP have been proposed to be APP-CT dependently regulated ^10–13^. In addition, some factors have been identified that modulate the subcellular localization of the APP-CT domain ^13–17^ as well as the molecules that are relevant for APP-CT-dependent DNA damage ^18,19^. However, APP-CT is also the target of a highly controversial and critical discussion, as many results, e.g. expression changes of its targets, could not be validated in independent laboratories ^20^. Whether this controversy is due to the use of different cell lines and expression constructs, or different experimental setups needs to be elucidated.

Promyelocytic leukemia nuclear bodies (PML-NBs) are multiprotein complexes with PML as the main building component ^21^. A diverse set of nuclear proteins have been identified as permanent or transient PML-NB-binding partners ^22^. PML-NBs are highly dynamic structures with respect to mobility, composition, architecture, and function ^23^. While the precise biochemical function(s) of PML-NBs have not been elucidated yet, they have been linked to many aspects of chromatin biology, including transcription, histone modification, repair and recombination, degradation, hence genome maintenance ^24^. Transcription of PML is strongly upregulated by interferons and p53 ^25^, causing a significant increase in the number and size of the bodies. Recent studies revealed an emerging role of PML-NBs as coregulatory structures of both type I and type II interferon responses ^26^. These data, along with the observation that a subset of PML-NB components (including PML, Daxx, and Sp100) can act as cellular restriction factors against viral attacks, strongly indicate that PML-NBs act in intrinsic antiviral defence mechanisms and innate immune responses. Within this work, we demonstrate that PML nuclear bodies interact with highly mobile APP-CT complexes and progressively form immobile large nuclear structures. Accordingly, each complex is able to mutually precipitate the other, and both reveal strong association up to complete co-localization. Our results further show that APP-CT co-localizes with PML in the human brain but not in cerebral organoids derived from wildtype (wt) iPS cells. In addition, plaque-rich areas of the diseased human cortex show a significant reduction in nuclei containing a single PML body compared to controls. We conclude that APP interaction with the nuclear PML complexes is age-dependent, which might be of relevance for AD pathophysiology. Based on these results, we suggest a new function of APP signalling in virus defence.

## Material and Methods

### Vector constructs

The plasmids encoding the fusion proteins (FE65-EGFP, FE65-mCherry, APP-CT-GFP, TIP60-EGFP, TIP60-HA, TIP60-BMP, EGFP-PML, PML-HA, PML-myc, p53-EGFP, Daxx-EGFP, H2A-mTurquoise, HIPK2-EGFP, HP1ß-EGFP, UBE2D2-mCherry, and WRN-EGFP) described in this paper, were generated using the In-Fusion^®^ HD cloning kit (Takara Bio) according to manufacturer’s instructions or were purchased (Addgene). Amplification and purification of the plasmids were done according to standard protocols.

### Cell culture, transfection, and immunofluorescence

Stem cells (iPS CD34) were cultured in StemFlex™ (Gibco) on 35 mm dishes, coated with Matrigel or Geltrex^®^ (Gibco) according to the manufacturer’s protocol, and split before reaching 70 % confluency. HEK293T cells were seeded and incubated in DMEM (Gibco) with 10 % heat inactivated FBS (Gibco), 1 % Penicillin/Streptomycin and 1 % L-glutamine (Gibco), to a confluency of 70 %. For overexpression assays, sterile precision cover glasses (1.5 H Marienfeld Superior) were placed into a 24-well cell culture plate (Sarstedt) and coated with 0.01 % poly-l-ornithine solution (Sigma Aldrich). The respective plasmids were transfected via the K4^®^ Transfection Kit (Biontex) according to manufacturer’s recommendations for 24-well culture plates. After 24 or 48 h, cells were briefly washed with DPBS (Gibco) and fixed in 4 % paraformaldehyde in PBS. Cover glasses were mounted with the Shandon™ Immu-Mount™ solution (Thermo Scientific) on glass slides and dried overnight at RT. For immunofluorescence staining, HEK293T cells were seeded in 8-well-μ-slide-ibi Treat (ibidi^®^, Martinsried, Germany) and transfected using calcium phosphate transfection after 24 h. Cells were fixed with Roti^®^-Histofix 4 % (4 % phosphate buffered formaldehyde solution; Roth, Karlsruhe, DE) for 20 min at 37 °C, and permeabilized and blocked with 5 % normal goat serum (NGS) in 0.3 % (w/v) Triton X-100/PBS for 30 min at RT. The cells were incubated with primary antibodies diluted in 1 % BSA/0.3 % Triton X- 100/DPBS (mouse anti-HA (BioLegend, 901501; 1:1000), mouse anti-myc (NEB/Cell Signaling, 2276; 1:1500) o/n at 4 °C. For the secondary antibody staining and the cell staining, goat-anti-mouse AF568 (Invitrogen, A11004, 1:1000) was used, together with HCS CellMask™ Deep Red Stain (ThermoFisher Scientific, H32721, 1:5000), Hoechst33342 (10 mg/ml in H2O, Applichem, A0741, 1:1000) in DPBS (1 % BSA, 0.3 % Triton) and incubated for 1 h at RT.

### Immunoprecipitation

HEK 293T cells were seeded in 10 cm dishes and co-transfection was performed 24 h after seeding. Whole cell extracts were prepared 24 h after transfection by scraping the cells from the dish with a cell scraper, washing the cell pellet in ice-cold PBS, extracting with 1 ml interaction buffer (50 mM Tris pH 8, 150 mM NaCl, 5 mM EDTA, 0.5% NP40, 1 mM DTT, 1 mM PMSF, 1x complete protease inhibitor cocktail), followed by sonication (15 s at 95 % amplitude) using a Sonopuls mini20 device (Bandelin, Berlin, Germany). The lysates were centrifuged (15,000 g, 15 min, 4 °C) and the supernatant was transferred to a new reaction tube. Input samples of the lysates were stored separately. Immunoprecipitation (IP) was carried out with the μMACS isolation kits for tagged proteins from Miltenyi Biotec (Bergisch-Gladbach, Germany). The eluates, as well as input samples of the lysates were subjected to SDS-PAGE and immunoblotting.

### Immunoblotting

Protein concentrations were determined using the Bio-Rad protein assay system (Bio-Rad Laboratories, Richmond, CA). Equal amounts of protein were resolved by SDS-PAGE using a 10 % acrylamide gel and subsequently transferred onto polyvinylidene difluoride (PVDF) membranes (Amersham Hybond, GE Healthcare) via the PerfectBlue™ tank electro blotter (Peqlab, Erlangen, Germany) with 350 mA for 90 min. To minimize unspecific binding, the membranes were blocked in 5 % (w/v) non-fat dried milk powder in TBST for 30 min at RT. Membranes were probed with primary antibodies against GFP (rabbit, polyclonal, 1:2000, Santa Cruz, sc-8334), HA tag (mouse, monoclonal, 1:1000, BioLegend, 901501) and p53 (mouse, monoclonal, 1:200, Novus Biologicals, NBP2-29419) diluted in blocking solution overnight at 4 °C. The membranes were washed three times with TBST, before they were incubated with HRP-conjugated secondary antibodies (1:10000, NXA931 and NA934, GE Healthcare Europe, Freiburg, DE) also diluted in blocking solution for 1 h at RT. Visualization of bound antibodies occurred via enhanced chemiluminescence (ECL) with the ECLplus Western Blotting Substrate from Pierce (Rockford, IL, USA) according to the manufacturer’s instructions. After incubation with the substrate, the detection of the generated signal was carried out with the ChemiDoc MP Imaging System (Bio-Rad Laboratories GmbH, Feldkirchen, Germany).

### Cerebral Organoids

Cerebral organoids were generated according to the protocol from Lancaster and Knoblich ^27^ with minor modifications, all media compositions remained unchanged. Briefly, at day 0, iPS CD34 positive cells were detached and harvested using TrypLE™ (ThermoFisher, Germany). Afterwards, DMEM/F12 was added to the detached cells and the cell number was calculated using a Neubauer chamber. Next, 9,000 cells/well were seeded into a 96-well ultra-low attachment plate (Corning) with a total amount of 150 μl hESC-medium (containing 4 ng/ml bFGF and 50 μM ROCK-Inhibitor) per well. On day 3, half of the media was exchanged with 150 μl of hESC-medium without bFGF and ROCK-Inhibitor. Subsequently (day 6), the embryoid bodies were transferred to a 24-well ultra-low attachment plate (Corning) with 500 μl Neural Induction (NI)-medium. Every day, half of the media was exchanged with 500 μl fresh NI-medium. On day 12, the embryoid bodies were embedded in droplets of Matrigel (Corning) and incubated for 25 min at 37 °C for Matrigel polymerization. Afterwards, the droplets were transferred to a 50 mm dish with differentiation medium without vitamin A (DM-A) medium for further incubation at 37 °C in a 5 % CO_2_ atmosphere. Four days later, the medium was changed to DM+A and the developing cerebral organoids (COs) were maintained at 37 °C with 5 % CO_2_ until experiments were performed.

### Histology and Immunohistochemistry

The COs were removed from the media, washed with PBS, and fixed with 4 % paraformaldehyde in PBS for 90 min at 4 °C. After washing with PBS, organoids were incubated in 30 % sucrose solution for cryoprotection at 4 °C overnight. The next day, the COs were embedded in a 1:1 mixture of 30 % sucrose and Tissue-Tek O.C.T. embedding medium (Science Services, SA62550-01), snap-frozen on dry-ice, and then stored at −80 °C until cryosectioning. Frozen COs were sliced into 15 μm sections using a cryostat (Leica CM3050S), mounted on SuperFrost™ slides (ThermoScientific™), and stored at −80 °C until further use.

For immunohistochemistry, COs and brain tissue sections were thawed for 2 min in PBS. To apply the biotin-avidin system used for the enhancement of fluorescence, the sections were first blocked with avidin for 10 min and, after washing twice with PBS for 4 min, blocked with biotin for 10 min. After washing twice with PBS for 4 min, sections were blocked and permeabilized in 0.1 % Triton X-100, 5 % goat serum in PBS for 1 h at RT, followed by incubation with primary antibodies in a humidified chamber overnight at 4 °C. Primary antibodies were diluted in 0.1 % Triton X-100 in PBS as follows: APP-CT (mouse, Millipore MAB343, 1:100), PML (rabbit, Novus Biologicals NB100-59787, 1:400), PML (mouse, Abcam ab96051, 1:200), TIP60 (mouse, Abcam Ab54277, 1:400), FE65 (mouse, Acris AM32556SU-N, 1:400), FE65 (rabbit, Santa Cruz sc-33155, 1:400), β-tubulin III (mouse, StemCell 01409, 1:100), p53 (mouse, Novus Biologicals NB200-103, 1:100). Sections were incubated with the biotinylated secondary antibody (goat anti-mouse IgG Biotin, Life Technologies B-2763, 1:100) diluted in 0.1 % Triton X-100 in PBS for 1 h at RT in a humidified chamber. Following washing twice with PBS for 4 min, sections were incubated with Avidin-TRITC (1:1000) and a non-biotinylated secondary antibody (donkey anti-rabbit FITC, Santa Cruz sc-2090, 1:100) diluted in 0.1 % Triton X-100 in PBS for 45 min in a humidified chamber protected from light at RT, and subsequently washed twice with PBS for 4 min.

In case of thioflavin-S counterstaining of amyloid plaques, sections were incubated with 0.1 % aqueous thioflavin-S solution, washed twice with PBS for 4 min, washed with 30 % ethanol followed by 50 % ethanol for 5 min each, and finally washed twice for 4 min with PBS.

For counterstaining of nuclei, DAPI solution (0.001 mg/ml) was added to the sections for 15 min while protected from light, then slides were washed twice with PBS for 4 min and mounted.

### Imaging and Tracking

Cells were either imaged after fixation and mounting (Shandon™ Immu Mount™ solution, Thermo Scientific) on glass slides or for life cell imaging directly using the integrated incubation chamber of the Leica (Mannheim, Germany) TCS SP8 microscope system (37 °C and 5 % CO_2_). Samples were imaged using a 63x water (1.2 NA) or 100x oil objective (1.4 NA). Fluorophores were excited with 405/488/514/561 nm laser lines performing a sequential scan beginning with the most red-shifted wavelength. Images were recorded into 1024×1024 images at a scan speed of 200 Hz with HyD detectors. Tile scans were imaged through the selection of 800 × 800 μm areas (5 × 5 tiles) in x- and y-direction. Additionally, z-stacks (n = 5) of 2 μm between each plane (8 μm in total) were recorded and merged via the maximum projection tool in the LASX-software (Leica, Mannheim, Germany). The fluorescence intensity curves were measured along the cell nucleus within a region of interest (ROI) and the chromatogram was normalized using the quantitative tools of the LASX-software tool (Leica, Mannheim, Germany). For the 3-dimensional imaging, several z-stacks (n=10) of 1 μm step size were recorded and the 3-dimensional image was generated using the LASX-software tool.

The track analyser of the Hyugens object tracker wizard was used to study the 3dimensional motion of the nuclear bodies of cells that were previously transfected with and without PML. Therefore, ROIs containing nuclear bodies only or background only were selected for the tuning of the detection filters via linear discrimination analysis (LDA) and the subsequently tracking of the nuclear bodies. The detection threshold was adjusted to measure objects with a positive generated score, computed by the software, to further discriminate against the background. The bodies were tracked within the cells over a time span of 300 s and the speed was calculated using the integrated software.

### STED Microscopy

For STED, GFP fusion proteins in fixed cells were labelled with Alexa Fluor 647-coupled GFP nanobodies (GFP-booster gb2AF647-50, Chromotek, Germany) at 1:100 dilution. Endogenous and overexpressed PML was immunofluorescently labelled with anti-PML antibody (rabbit, ABD-030, Jena Bioscience, Germany, 1:500), followed by secondary antibody coupled with STAR 580 STED dye (goat-anti-rabbit, ST580-1002-500UG, Abberior, Göttingen, Germany, 1:100). Stained cells were embedded in ProlongGold with DAPI (Thermo Fisher Scientific, Germany) and covered with 12 mm round cover glasses (Thickness 0.17 ± 0.01 mm). Gated STED images were acquired on a Leica TCS SP8 STED microscope equipped with a 100x oil objective (HC PL APO CS2 100x/1.40 Oil) according to protocols established for nuclear bodies by Okada and Nakagawa ^28^. Pixel size in STED acquisition was applied automatically in LAS-X software (Leica, Mannheim, Germany) for the most red-shifted dye (AF 647), usually resulting in a pixel size of less than 20□□ū20 nm. STED beam alignment was performed before each imaging session between the pulsed white light laser and the 592 nm depletion laser. DAPI, Alexa Fluor 488, Star 580 and Alexa Fluor 647 were excited with laser lines 405 nm, 488 nm, 580 nm and 635 nm of the white light laser, respectively. Emission was captured through band pass settings 430-470 nm, 505-550 nm, 590-620 nm and 648-720 nm, respectively. Depletion of STAR 580 and AF 647 was performed with the 775 nm depletion laser. The power of the depletion laser was optimized for each dye to obtain highest resolution while minimizing bleaching. Imaging conditions were fine-tuned on several cells before application of the optimized settings for final images. Each dye was imaged in sequential scans to avoid spectral overlaps. While hybrid detector gain was set to 100 %, excitation laser intensity was set such to prevent pixel saturation. Images were obtained using a pixel dwell time of 100 ns. Photon time gating was employed by collecting lifetimes between 0.3 and 6.0 ns. To compensate for inevitable signal intensity loss during STED acquisition, the excitation laser power was set 3- to 5-fold higher than in conventional confocal mode. When using STED channels, the pinhole was set 1.0 Airy Units. In non-STED channels the pinhole was set to 0.49 Airy Units to allow for sub-Airy super-resolution confocal microscopy according to the HyVolution II mode of the Leica SP8 microscope system. All images were deconvolved with Huygens Professional Software (Scientific Volume Imaging B.V., Hilversum, The Netherlands) using the deconvolution presettings in Huygens software applying Classic Maximum Likelihood Estimation (CMLE) algorithms.

### Cell Profiler Data Analysis

Brain tissue slides acquired from patients with Alzheimer’s disease in different stages of severity, were stained with Thioflavin, DAPI and anti-PML antibodies (as described before). The slides were imaged using the Leica TCS SP8 confocal microscope system with a 100x oil objective (HC PL APO CS2 100x/1.40 Oil) and tile scans (10 x 10 tiles) containing amyloidogenic plaques were recorded. Additionally, several plaque-free regions were captured as control (n = 9 from 3 individuals). PML bodies and cell nuclei were identified and counted using the CellProfiler™ Software. In the tile scans, rectangular areas around the plaques were defined as ‘plaque near’ (n = 31 from 8 individuals) and the surrounding area as ‘plaque distant’ (n = 28 from 7 individuals), not including cells close to the edge of the tile scans. The acquired data from CellProfiler™ were exported to Excel, sorted in those two groups and compared regarding the amount of PML bodies inside the cells and the percentage of cells containing those aggregates.

## Results

### The C-terminal domain of the amyloid precursor protein induces gene expression active aggregates with donut-like shape in a variety of cells

Starting point of this study was the idea to validate or discard the potential signal transduction pathway caused by the amyloid precursor protein C-terminal domain (APP-CT), which has been controversially discussed over the last years ^11,20,29–32^. The stumbling block of APP signalling is the high instability of its cleaved C-terminal fragment, its fast degradation (on the other hand its stabilization by interacting proteins), and the use of non-appropriate models ^*30,33–36*^. In order to overcome at least some of those hurdles, we initiated our study by the analysis of potential nuclear APP signalling in various cell lines including primary neurons (Figure 1). APP-CT forms nuclear aggregates with two other proteins, which has been reported by several groups in overexpression systems^12,29^. These are the histone acetyltransferase TIP60 and FE65, an adapter protein with strong APP-CT binding capacity. The sequestered APP-CT subsequently establishes a triple nuclear complex with TIP60 and FE65, although the expression of FE65/TIP60 alone (without APP-CT) is sufficient for the phenotypical generation of these aggregates (which might only be the case using ectopic expression) ^9^. Confirmatively, many different cell lines including cancer cells, fibroblasts and primary neurons revealed the typical dot-like phenotype upon co-expression of FE65/TIP60 (Figure 1A). All cell types analysed here exhibited the same phenotypes - from cells with many tiny aggregates (Figure 1B, arrow) to those with a few large speckle-like structures (Figure 1B, arrowhead; an overview image is also shown in Supplementary Figure S1). As also live cell imaging validated an organization resembling a highly dynamic circular structure, it was initially speculated that the aggregates correspond to intranuclear lipid droplets. However, electron microscopy analysis (Figure 1C; Supplement Figure S2) as well as CellMask™ membrane stain (Supplement Figure S3) argued against this hypothesis and rather revealed a donut-like shape of the intranuclear aggregates with an electron-dense border and an electron-poor centre.

**Figure 1.**
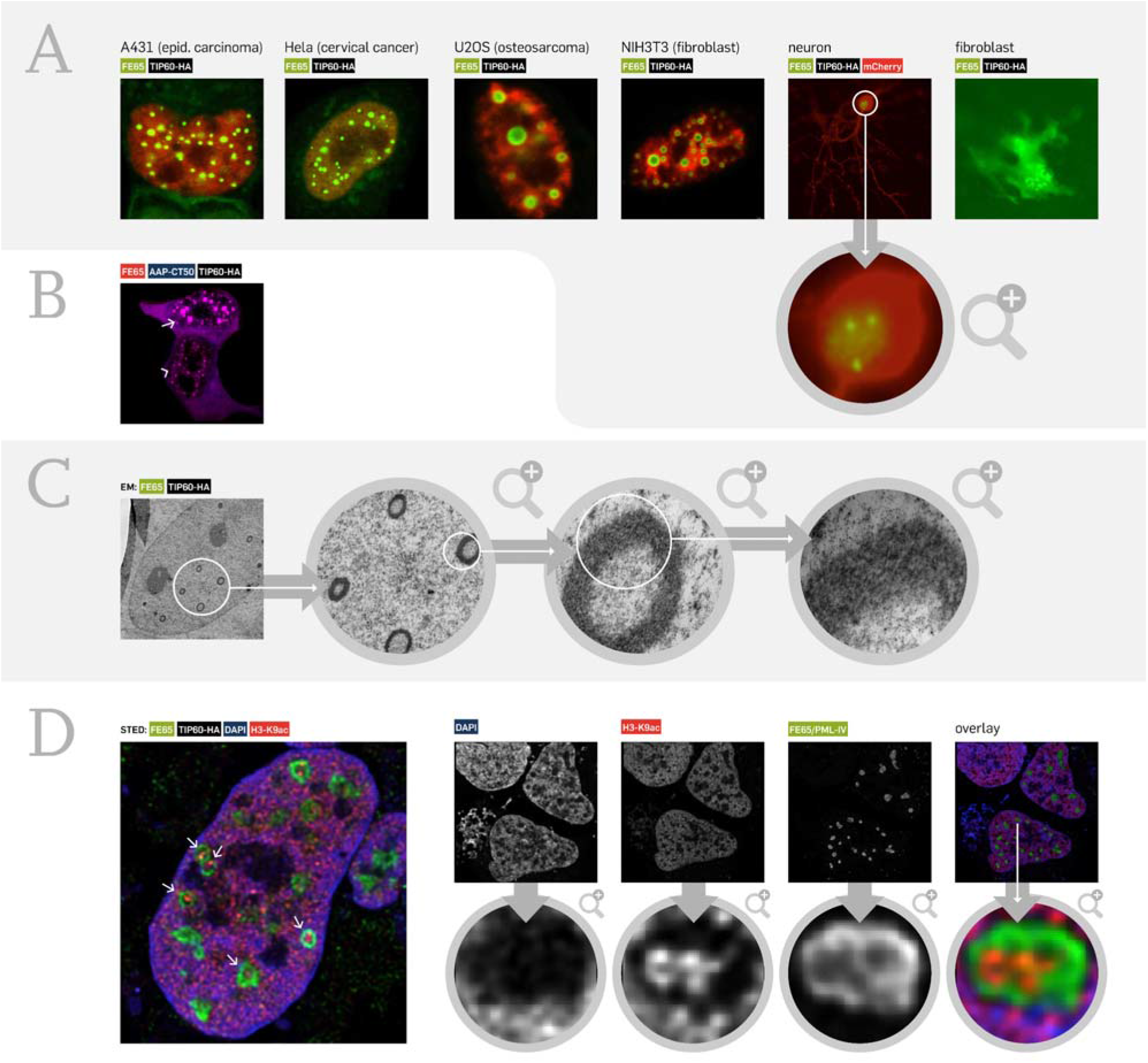
Dynamic nuclear APP-CT dependent aggregates are present in various cells, lack a membrane coating and are transcriptionally active. **(A)** Upon co-expression of FE65-mCherry/TIP60-EGFP, nuclear aggregates are generated in various cell lines including neurons. Transfected vectors with respective fluorophore (EGFP, mCherry) are indicated, TIP60 was co-transfected w/o fluorophore. **(B)** Every cell type used in **A** demonstrated cells with many tiny (arrow) or few large spheres (arrowhead) (or transition states) supporting the hypothesis of sphere fusion over time (blue, DAPI counterstain of nuclei; an overview image is also given in Supplement Figure S1). **(C)** Transmission electron microscopy of FE65/TIP60 transfected cells revealed an electron-dense ring structure (additional image in Figure S2). However, there was no evidence for a membrane sheath. These results were also confirmed by a CellMask staining (Supplement Figure S3). **(D)** High-resolution STED imaging revealed that the inner core of the aggregates is positive for anti K9 acetylation histone 3 antibody staining (red) supporting the hypothesis of active gene expression within the aggregates.

According to present literature^10–12^, a potential function of the APP-dependent nuclear aggregates is the modulation of gene expression in dependence of yet unknown stimuli. In order to test this hypothesis, transfected cells were fixed and stained with an anti-Histone3-K9ac antibody recognizing transcriptional active loci. Indeed, high resolution STED indicated active gene expression (positive staining in red) within large ring-like structures (Figure 1D, arrows; Supplement Figure S4). Thus, these data suggest that APP contributes to the formation of a nuclear membrane-free complex with impact on DNA-associated processes.

### The nuclear APP-CT complex associates to two tumour suppressor proteins

Stimulated by these promising results, we subsequently aimed to identify the protein composition of these structures in more detail. However, initial attempts to isolate the complexes by precipitation, nuclear enrichment, or filtration techniques all failed or did not yield a sufficient purity. This might potentially result from a strong association of the aggregates with the rather sticky DNA molecules. As an alternative approach, we extracted a set of proteins from the literature revealing a nuclear phenotype similar to the APP-CT-dependent aggregates, which resulted in a list of the following proteins: Daxx, H2A, HIPK2, HP1ß, p53, PML, UBE2D2, and WRN. For all of these proteins, vectors encoding the respective candidate DNA sequence fused to a fluorescent protein cassette were cloned and co-transfected with expression constructs for the APP-CT complex (APP-CT/FE65/TIP60). The vast majority of our candidates showed no co-localization; however, there were two exceptions. One protein, which revealed a co-localization, has been described before to be part of nuclear bodies ^37^ and to bind to BLM, which is an earlier described interactor ^*9,38*^, as well as to reveal an unstable interaction to TIP60 ^*39*^: the tumour suppressor protein p53. Indeed, co-expression of p53-EGFP, FE65-mCherry and TIP60-BFP demonstrated a strong co-localization in nuclear aggregates (Figure 2A). This could be validated by profiling of the individual fluorescence intensities (Figure 2B), all three fluorescent signals revealed the same intensity course along the dotted line with peaks at position I, II, and III (Figure 2B). Omitting FE65 co-expression demonstrated p53 co-localization to TIP60 speckles alone as well (Figure 2C), which is in line with earlier results ^39^. To further confirm the interaction of p53 with APP-CT-depending complexes, co-immunoprecipitation assays (co-IPs) were performed (Figure 2D). Sample conditions were selected as indicated in the input blot (Figure 2D, left). Co-IP (Figure 2D, right) using anti-HA tag antibody (TIP60-HA) revealed precipitation of FE65 (as expected, Figure 2D, white arrow) and of p53 (red arrow), which validated its participation in the complex. The TIP60/p53 interaction (independent of FE65, in agreement to Figure 2C) could be confirmed by precipitating TIP60 via a GFP tag antibody (white arrowhead). Here, in addition to the p53-EGFP signal, the presence of co-precipitated endogenous p53 was observed (3rd land, red arrowhead). Endogenous p53 is also detectable in conditions with TIP60 co-expression (e.g. lane 7, blue arrow) suggesting an interaction of the (endogenous) tumour suppressor protein p53 with the histone acetyltransferase TIP60 in nuclear aggregates.

**Figure 2.**
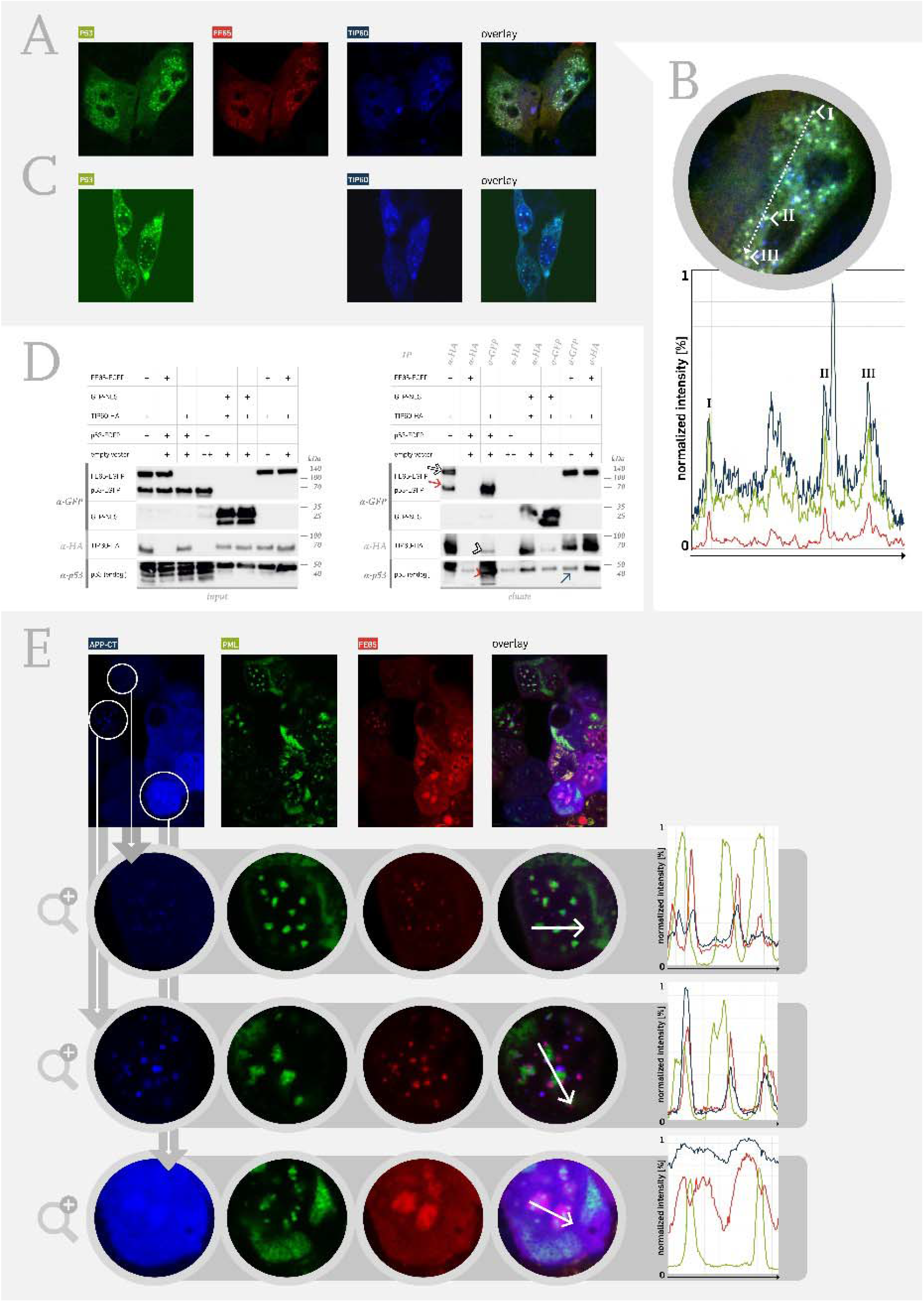
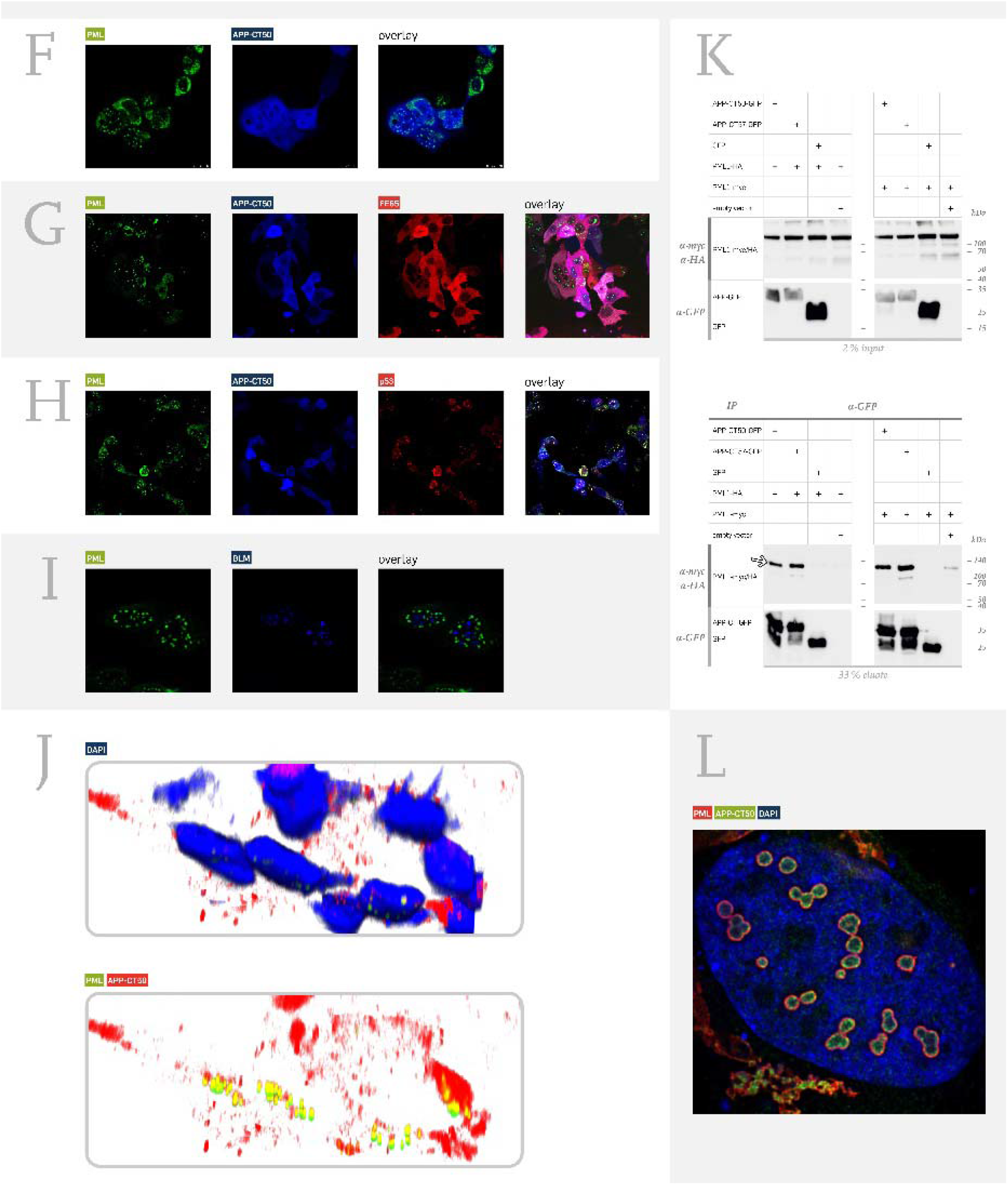
The nuclear APP-CT complex associates to p53 and PML. **(A)** The tumour suppressor protein p53 revealed the same nuclear dot-like structure as the co-transfected components FE65 and TIP60. Fluorescently labelled proteins are indicated in the Figure by the respective colour. **(B)** Tracking of the fluorescence intensity along the indicated arrow revealed peak intensities of each component (green: p53, red: FE65, blue: TIP60) supporting the association within one complex. **(C)** P53 is also co-localized to TIP60 independent of FE65. **(D)** P53 interaction with the APP-CT complex (FE65-EGFP/ TIP60-HA co-transfected to p53-EGFP vs. control (GFP-NLS, nuclear localization sequence)) was validated by co-immunoprecipitation (above: input blot, below: elution blot). IP using anti-HA tag antibody (against TIP60-HA) revealed precipitation of FE65 (white arrow) as well as of p53 (red arrow). Respective controls did not show a co-precipitation. TIP60 precipitation also occurred using anti-GFP as bait in well agreement to results obtained in part C (white arrowhead). Notably, high levels of endogenous p53 co-eluted in the same condition (red arrowhead), whereas a moderate endogenous p53 signal was observable in control conditions (red arrow). **(E)** A second tumour suppressor protein was identified to associate with the APP-CT complex: the promyelocytic leukemia protein PML. Different phenotypes of association were observed, e.g. one or two APP-CT/FE65 dots (TIP60 was co-transfected w/o fluorophore) associated with a single PML aggregate (first and second zoom-in row, compare fluorescence intensities). Alternatively, large APP-CT/FE65 complexes with enclosed PML-dots were found (third row). **(F)** A direct co-localization of APP-CT with PML was only observed to some extent (cells with nuclear aggregates), but most cells revealed a uniform staining pattern of APP-CT50 and PML in the cytosol and nucleus. **(G)** Additional coexpression of FE65 enriched the aggregation of PML in the nucleus, but a strong colocalization to APP-CT/FE65 was not observable in the imaging study. **(H)** p53 is part of the PML aggregates in the nucleus that also contain APP-CT. **(I)** Notably, the DNA helicase BLM, which was identified as binding protein in the APP-CT complex, is not co-localized with the PML aggregates. **(J)** Confocal 3D imaging validated the co-localization of PML and APP-CT in the nucleus (FE65/TIP60 were co-transfected). **(K)** Interaction of APP-CT with PML was shown using co-immunoprecipitation assay. Precipitation using anti-GFP antibody revealed detection of APP-CT-GFP (as expected) as well as PML (PML1 isoform was used, white arrow). This was true for two different APP-CT isoforms (APP-CT50 and CT57), whereas control conditions revealed no unspecific co-precipitation. Results were the same for two different PML1 tags: in the left panel of blots PML1-HA was used, whereas PML1-myc was used in the right panel. **(L)** High-resolution STED imaging specified the localization of APP-CT within the PML bodies. Different phenotypes were evident, either with a uniform localization within the bodies or with APP-CT signal at the inside wall of the PML body (red: PML, green: APP-CT).

The second candidate that aroused our interest was a tumour suppressor protein as well. It is known for its assembly in nuclear bodies amongst chromatin and its role in apoptosis, genome stability, and cell division: the promyelocytic leukemia protein (PML) ^21^. Coexpression of PML-GFP, BFP-APP-CT, FE65-mCherry and TIP60 (w/o fluorophore) revealed a close association of PML bodies to one or two APP-CT aggregates (Figure 2E, zoom 1), which is also confirmed by the fluorescence intensity line scan given in Figure 2E on the right. Indeed, the green PML peak is accompanied by two peaks of APP-CT (blue) and FE65 (red). Other cells of the same condition (zoom 2) demonstrated association of three APP-CT complexes, whereas another phenotype (zoom 3) demonstrated large APP-CT complexes with incorporated PML bodies. Collectively, the high dynamics of the nuclear complexes was compelling and pointed to a spatial-temporally highly organized mechanism. Direct co-localization of PML with the APP-CT fragment (w/o FE65/TIP60 co-expression) was also observed in a minor percentage of transfected cells (not shown), however, the main phenotype revealed a uniform APP-CT signal without accumulation in nuclear aggregates (Figure 2F). Nevertheless, cytosolic PML might also be bound to APP-CT. Additional expression of FE65 did not change the main phenotype of uniformly localized APP-CT (Figure 2G). Omitting TIP60 co-expression caused the generation of a PML/APP-CT/p53 complex (Figure 2H). In contrast, the DNA helicase BLM (ectopically expressed), which has been described to bind to the APP-CT complex, is not co-localized to PML (Figure 2I) demonstrating that not every highly expressed nuclear protein is in complex with PML. A more detailed analysis, utilizing 3D confocal imaging, revealed that APP-CT and PML are associated with each other in all nuclear aggregates (Figure 2J). Finally, the PML/APP-CT interaction was confirmed by a co-immunoprecipitation assay (Figure 2K). Co-expression of APP-CT-GFP with either PML1-HA or PML1-myc demonstrated precipitation of PML (detection via anti-myc or anti-HA antibody, Figure 2K, white arrow) upon anti-GFP IP (independent experiment given in Supplement Figure S5). This was true for two different APP-CT isoforms with 50 and 57 aa in length that are existent upon γ-secretase-mediated APP cleavage. High-resolution STED imaging further specified the co-localization of PML and APP-CT within the PML bodies. APP-CT was either uniformly distributed or concentrated at the inner wall of the nuclear body (Figure 2L). These results suggest that upon translocation of APP-CT into the nucleus, there is a regulated interaction between the APP-CT/FE65/TIP60 complex, p53 and PML bodies ^40^.

### APP-CT depending complexes drive PML complex generation that are also present in the aged human brain

To further investigate this interaction, live cell imaging experiments were performed in HEK293 cells with ectopic expression of the aggregate components. Expression of FE65-GFP, TIP60 (w/o fluorophore) revealed highly dynamic ring-like structures moving throughout the nucleoplasm (Figure 3A, arrowhead, confocal image, structures coloured according to z-level). To investigate the influence of PML in these dynamics, the complexes (based on the FE65-mCherry signal) were tracked in APP-CT50/FE65/TIP60-transfected cells with vs. without PML co-expression (Figure 3B). The velocity of the nuclear structures in the presence of PML (Figure 3B, diagram 1, red graph) was significantly reduced compared to the condition without PML (black graph). Moreover, the average distance from the track origin was significantly less in PML co-expressing cells (Figure 3B, diagram 2). The mean speed was 0.38 in PML vs. 0.76 μm/s in non-PML co-expressing cells (Figure 3B, diagram 3). These results suggest a mutual trapping function of APP-CT aggregates and PML, for which Figure 3C shows the typical phenotype of up to three APP-CT aggregates (red) bound to a single PML body (green). In order to understand the conditions for aggregation in more detail, we monitored transfected cells over time (Figure 3D). Pure PML expression (upper row) revealed a homogenous distribution of the protein within the whole cell (cytoplasm and nucleus). Co-transfection of APP-CT (blue) and FE65 (red) caused late aggregate formation mostly after 96 h (middle row), whereas expression of PML/FE65/APP-CT/TIP60 demonstrated nuclear body formation already after 24 h (below row). After 48 h, some cells revealed formation of super-aggregates within the nucleus (white arrow). We subsequently aimed to investigate whether these complexes are present in the human brain as well (Figure 3E). Human hippocampal frozen brain samples from 15 AD patients (with different Braak stages) were used to study the potential co-localization of APP-CT and PML. Adaption of the immunofluorescence protocol (compare methods part) was crucial for successful staining and indeed, we observed a strong co-localization of PML and APP-CT in the human brain (additional images as well as negative control omitting primary APP-CT antibody are given in Supplementary Figure S6). Tracking of the fluorescence intensity along the white arrow (including three aggregates) validated the strong co-localization. Stimulated by these encouraging results, we also investigated the localization of FE65 in the same manner (Figure 3F) and demonstrated that FE65 co-localized with PML in the human brain as well. Both co-localizations (APP-CT and FE65 with PML) were evident in the brains of AD patients with different Braak stages. As all samples were obtained from individuals older than 65 years, we next aimed to study whether this phenotype is age-dependent. To study this question in a human 3D model, we differentiated iPS cells to cerebral organoids (Figure 3G). Embryoid bodies (EBs) were generated followed by Matrigel embedding to differentiate 3D tissue-specific features of the human brain. In contrast to the aged human brain, immunofluorescence analysis demonstrated no co-localization of APP-CT with PML in cerebral organoid sections. These data suggest that the nuclear APP-CT/FE65/PML aggregation occurs in the human Alzheimer brain and might be an age-dependent effect.

**Figure 3.**
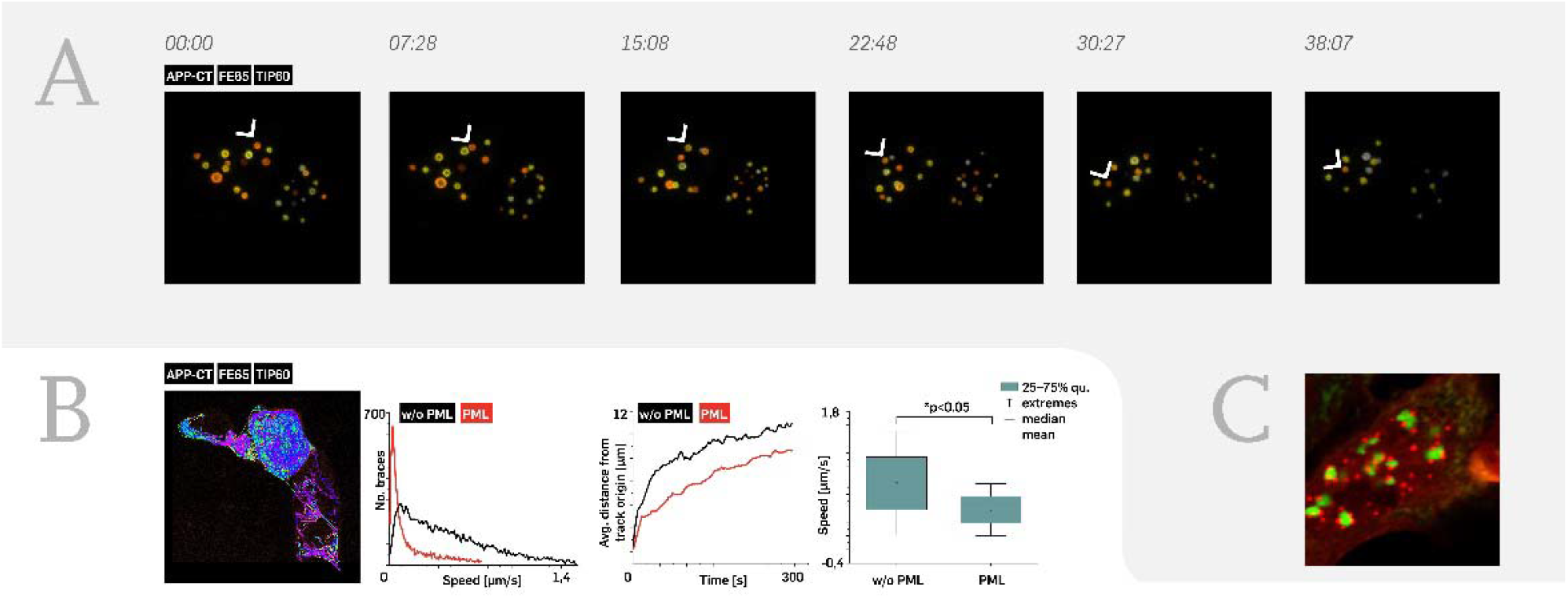
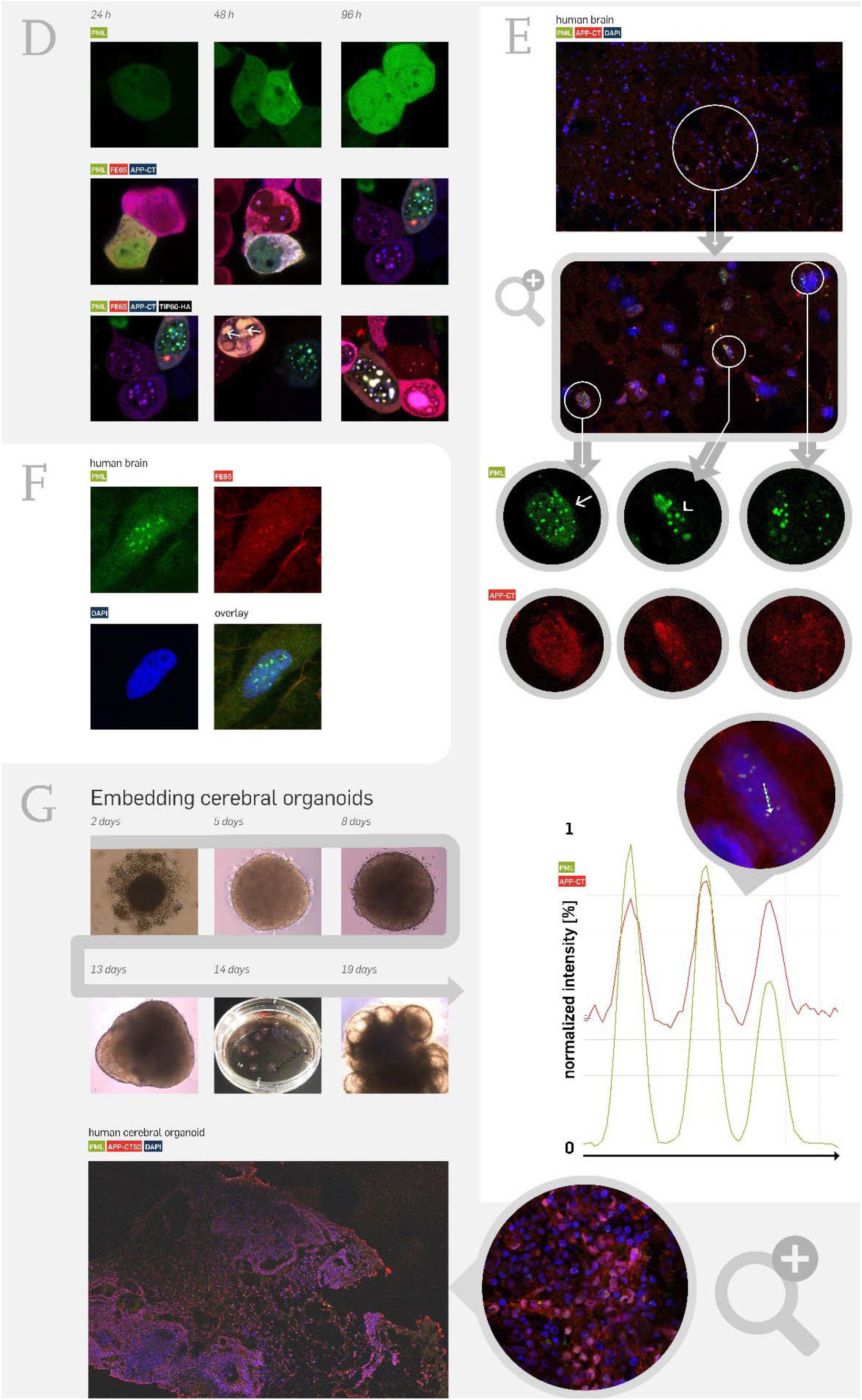
Highly mobile APP-CT-depending complexes that are also present in the aged human brain drive PML complex generation. **(A)** Expression of APP-CT/ FE65/TIP60 in HEK293 cells reveals a highly mobile complex moving three-dimensionally in the cellular nucleus. Ring-like structures were coloured (orange to yellow) according to z-level (confocal microscopy). The indicated structure (white arrowhead) revealed time-dependent movement. **(B)** Movement of the individual aggregates was tracked using Huygens object tracker software. Transfection in HEK293 cells included APP-CT/FE65-mCherry/TIP60 with and without GFP-PML co-expression. The FE65-mCherry signal was used for tracking, revealing lower speed in cells co-expressing PML (first diagram). In addition, the distance from the track origin (at time point 0) was analysed. Co-expression of PML revealed significantly lower distances pointing to mutual trapping of both complexes. The mean speed was 0.38 in PML vs. 0.76 μm/s in non-PML co-expressing cells (last diagram). **(C)** This part reveals a representative image demonstrating the complex generation of APP-CT/FE65/TIP60 (red) and PML (green aggregates). **(D)** Time-dependent generation of nuclear APP-CT/PML aggregates. EGFP-PML expression revealed a uniform distribution within the nucleus and cytosol (first row). Co-expression of FE65-mCherry and APP-CT caused initial aggregate formation after 48 h (middle row). Additional expression of TIP60 (last row) showed early generation of nuclear aggregates after 24 h. 48 h after transfection large nuclear aggregates were observed (white arrow). **(E)** PML/APP-CT colocalization was studied in human brain sections. In total, 15 human hippocampal sections were analysed (different Braak stages). Confocal tile-scan imaging (5 z-stacks, then fused by maximum projection algorithm) revealed strong co-localization of PML (green) with APP-CT (red). As in cell culture experiments, nuclei containing many small aggregates (arrow) as well as nuclei with larger aggregates (arrowhead) were evident. Co-localization is further shown by intensity tracking of both fluorescent channels in the diagram for a nucleus along the dotted white arrow. **(F)** Similarly, co-staining of PML and FE65, which confirmed the association of both proteins in the nuclei of the human brain, was performed. **(G)** In order to address the question whether co-localization also occurs in non-aged tissue, we differentiated human cerebral organoids from induced pluripotent stem cells. Embryonic bodies were embedded in Matrigel at day 11 followed by neuronal induction to generate organoids, which were analysed after 30 days in culture (seeding at day 0). Staining of cryosections failed to demonstrate co-localization of APP-CT and PML.

### Reduction of PML bodies occurs in human hippocampal brain areas with high plaque load

In order to study a potential pathophysiological relevance, we examined human brain tissue in more detail (Figure 4). As the human hippocampus belongs to those brain areas revealing early pathological AD features, we studied the extent of PML bodies in the Cornu Ammonis areas 1 or 3 (CA1, CA3). CA regions were not evident for all human brain samples due to different quality (different post-mortem times, preparation artefacts), thus we limited our analysis to CA1 or CA3 areas, which were distinctly assignable, e.g. Figure 4A corresponds to a sample with CA1 assignment, but not CA3. Haematoxylin Eosin (HE) staining of human hippocampal frozen sections (samples from 15 AD individuals with different Braak stages; due to the limited number of cases we did not analyse CA1 and CA3 separately) was used to identify the specific areas. Parallel sections (from the same individual) were used for low-resolution tile-scale imaging (DAPI channel, Figure 4A) to allocate hippocampal areas according to HE staining. For subsequent detailed analysis, confocal tile-scan imaging was used including 5×5 tiles and 5 z-stacks (Figure 4B). Afterwards, maximum projection algorithm was applied resulting in high-resolution imaging enabling identification of the number of PML bodies in each nucleus over an area of 800 x 800 μm within CA1 or CA3 (Figure 4C). This imaging pipeline was further extended to identify areas of high plaque load (within CA1 or CA3) using Thioflavin co-staining (Figure 4D). In total, we scanned 68 areas using this approach. The decision to use the PML signal (instead of APP-CT or FE65) for the subsequent quantitative analysis was made on the basis of best signal to noise values (compare Figure 3E and F). All tile-scans were subsequently processed using CellProfiler™ software in order to identify and extract nuclei (Figure 4E, upper row) and to detect and count the number of PML bodies within the extracted nuclei (Figure 4E, bottom row). Data analysis of all cells containing between 1 to 5 PML bodies compared to all cells counted (cells without any PML body, cells with more than 5 PML bodies), revealed a significant (p<0.05) reduced percentage of PML positive nuclei in areas with high plaque load (Figure 4F, plaque) compared to plaque free areas (Figure 4F, no plaque). Tile-scans from individuals without any neurodegenerative pathology (controls) again demonstrated higher percentage of PML positive nuclei compared to the “no plaque” group (p<0.05). More detailed analyses demonstrated that the observed significance is particularly caused by cells containing only one or two PML bodies per nucleus (Figure 4G, *p<0.05, **p<10^-5^). Taken together, these results suggest that human brain areas with high APP processing, thus causing a high plaque load, correlate with a significant reduction of the number of PML bodies per nucleus. Thus, APP signalling to the nucleus might regulate the generation and fusion of these aggregates which contain APP-CT, FE65, p53, and PML.

**Figure 4.**
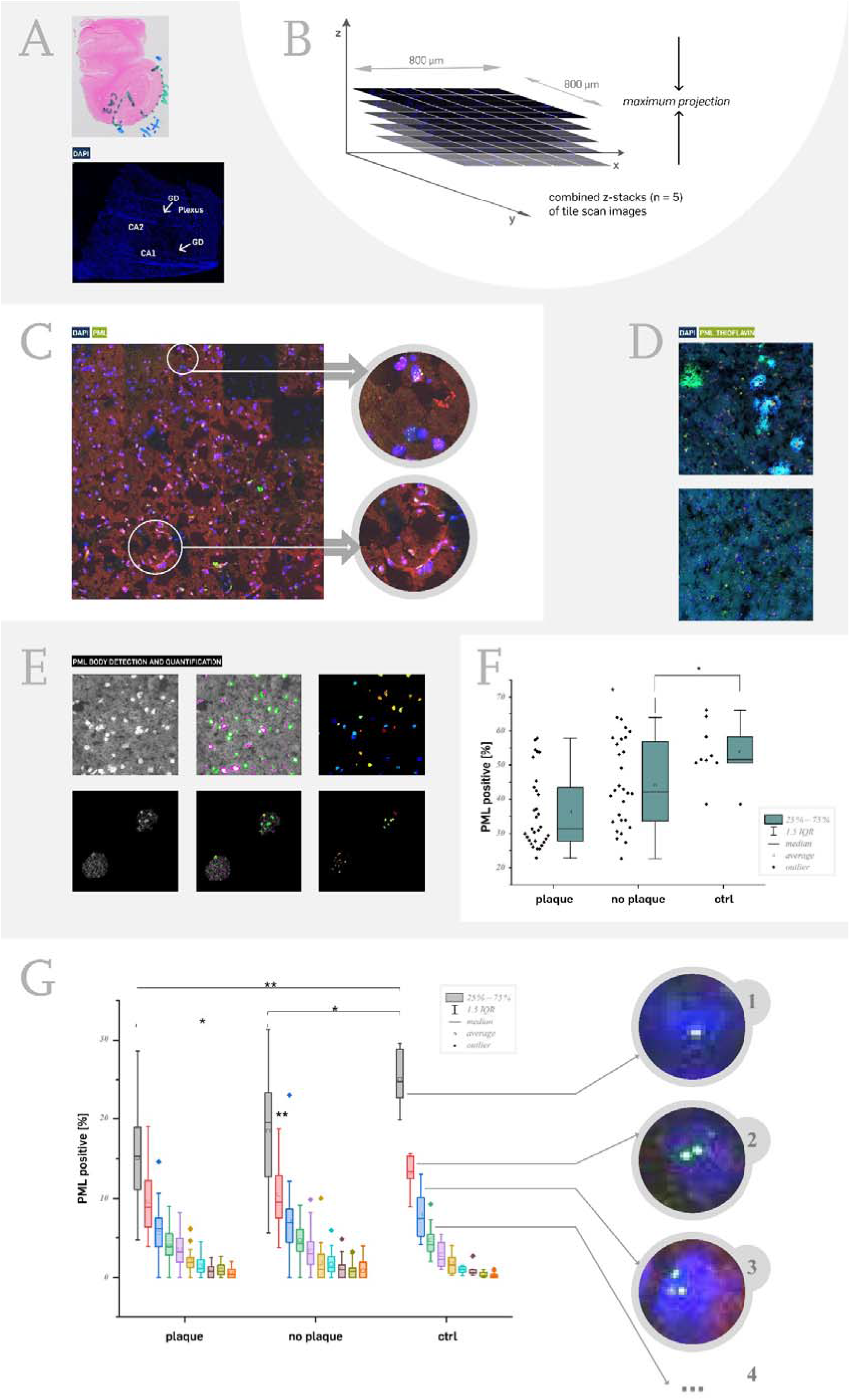
Significant enrichment of PML bodies in human brain areas with high plaque load. **(A)** Haemotoxylin eosin (HE) staining of human hippocampal sections was used to define Cornu Ammonis (CA) 1-3, Gyrus dentatus (GD), and Plexus areal. Parallel sections were used for PML immunofluorescence staining. DAPI co-staining was used for low-resolution tile-scan imaging and allocation of specific CA areas according to the initial HE staining. **(B)** Hippocampal CA1 or CA3 areas were used for high-resolution tile scan imaging. A single scan included 5×5 images with 5 z-stacks, which were subsequently combined using maximum projection. **(C)** A representative high-resolution (100x objective) confocal tile-scan image demonstrates identification of PML bodies in DAPI counter-stained nuclei. **(D)** Tile-scan imaging was then established with Thioflavin co-staining in order to identify areas with high plaque load in the human hippocampus (CA1 or CA3). **(E)** CellProfiler™ software was used to automatically annotate and extract nuclei (DAPI channel in grey scale) from the image (upper row). Subsequently, PML body identification and quantification was done in the extracted nuclei. **(F)** All nuclei containing 1 to 5 PML bodies were used to determine the percentage of PML positive nuclei (y axis). Hippocampal areas with high plaque load (plaque, average = 36.2 %) revealed a significant lower percentage of PML positive nuclei compared to areas without plaque (no plaque, average = 44.1 %) (p<0.05; every dot (left to each bar) indicates a single tile-scan experiment; for areas with high plaque load 31 tile-scans were analysed and quantified, 28 for no plaque, 9 for control). Tile-scans of control sections (individuals without any plaque pathology, average = 53.8 %) revealed again higher percentage of PML positive nuclei compared to “no plaque” areas (p<0.05). **(G)** Detailed analysis of cells containing one up to ten PML bodies per nucleus revealed significant differences. Nuclei with a single PML body were underrepresented (p<0.05) in areas with high plaque load vs. areas without plaques. Difference to control tile scans were highly significant (p<10^-5^). In addition, nuclei with two PML bodies (red bars) also revealed highly significant differences to the control. Representative images for nuclei containing one, two, or three PML bodies are given.

## Discussion

The amyloid precursor protein is a ubiquitously expressed protein, thus it is not surprising that many different cell types are capable to induce APP-CT signalling and to set up the nuclear aggregates. FE65 was initially reported to be brain tissue-specific, and indeed, expression analysis suggested FE65 to be a neuronal protein ^41–43^. However, an alternatively spliced FE65 isoform was found exclusively expressed in non-neuronal cells ^44^ and a role as transcriptional co-regulator was reported in breast cancer cells ^45^. TIP60, the so far described third player in the complex, is also ubiquitously expressed with the highest amounts in testis and placenta. This protein plays a general role in the acetylation of histones decreasing their binding affinity to the negatively charged DNA ^46^. Thus, we conclude that APP-CT/FE65 signalling is a ubiquitous pathway with a preference in neuronal cells due to neuron-specific FE65 splicing.

Notably, APP-CT dissociation and nuclear translocation was described to predominantly occur through the amyloidogenic processing pathway ^31^, which points to a pathophysiological function of APP-CT and suggests that the same is true for the role of APP-CT in the nucleus. This hypothesis is further supported by findings in APP-CT overexpressing mice revealing neuronal network abnormalities ^47^. Unfortunately, it is unknown whether these mice exhibit the nuclear APP-CT-dependent complex phenotype and whether the observed changes are based on gene expression changes, which has not been studied in these models so far. However, that APP-CT has an impact on gene expression has been suggested by different groups ^10–13^, and our findings of positive histone 3 K9 acetylation in the core of the APP-CT aggregates further support this hypothesis. The donut-like structure of the nuclear aggregates fits to this function as well, assuming that DNA is incorporated in (or associated to) the spherical aggregates as already shown for cells in G2 phase ^48^. Thus, APP-CT aggregates might correspond to DNA incubation containers modifying DNA in a way to change expression. The identification of reliable gene targets, which are differentially expressed, is so far hampered by the lack of suitable models. Until now, overexpression models have been used to identify potential (critically discussed) targets of the APP-CT complex ^11^. However, gene expression is just one suggested model, thus APP-CT complexes might also be relevant for any other nucleic acid mechanisms. The high mobility and fusion of the nuclear complexes, which generate larger and larger aggregates over time ^9^, provoked the presence of a vesicle-like, membranous structure, which is, however, not supported by our experiments. Alternatively, the APP-CT aggregates might correspond to condensed liquid droplets, which are slightly denser than the bulk intra-nuclear fluid ^49^.

In order to better understand the nuclear APP-CT-depending aggregates, we aimed to identify new protein components and were indeed able to validate the two tumour suppressor proteins p53 and PML as additional components. According to our results in HEK293 cells, this interplay is strongly driven by the APP-CT nuclear translocation, as pure PML expression revealed a rather homogenous cellular distribution within cytosol and nucleus. The complex generation follows a temporally organized scheme with generation of small aggregates at an early phase and few large nuclear complexes at later time points. As these aggregates were evident in the human brain of aged patients but not in cerebral organoids, we conclude that the APP-CT nuclear signalling is age-dependent and potentially of relevance for the pathophysiology of Alzheimer’s disease. Staining of APP-CT and FE65 was only successful using a sophisticated protocol with defined order of antibody incubation, meaning that incubation with the secondary antibody for PML detection precluded the identification of APP-CT or FE65 in the complex. In contrast, usage of the secondary antibody as the last step (after application of the Avidin-TRITC complex to identify APP-CT of FE65) was successful to reveal co-localization. Thus, accessibility of the epitopes of APP-CT or FE65 was presumably masked by the secondary antibody used for PML staining, suggesting a localization of APP-CT and FE65 within PML spherical aggregates, which is in good agreement to other reports ^48^.

Relevance of APP-CT/FE65/PML aggregates for the pathophysiology of AD was finally demonstrated by tile-scan imaging of hippocampal CA1 and CA3 areas. Due to the limited availability of high-quality frozen human hippocampal sections, our study had to be reduced to 15 AD cases with different Braak stages, in that we analysed 68 tile-scan areas in total. Although we were facing these limitations, our data analysis revealed highly significant results demonstrating a reduction in the number of PML bodies in nuclei close to AD relevant hot spots with high plaque load. Assuming that these areas also correspond to elevated APP cleavage, we conclude that APP nuclear signalling involving the adapter protein FE65 is also correlated to AD pathology. These results correlate well with our findings in cell culture. Indeed, expression of APP-CT, FE65, TIP60 and PML in HEK293T cells caused a time-dependent fusion of the nuclear aggregates. Thus, APP-driven fusion of PML aggregates seems to occur in the AD brain as well. Certainly, further studies are pivotal to understand the consequences of APP to PML body generation, fusion, and, in particular, their impact on neurodegeneration. Although, our present knowledge on PML body function is limited, highly interesting hypotheses in the context of neurodegeneration can be derived (Figure 5). Based on our results and the current literature, we suggest a new model for the role of nuclear APP signalling in neurodegeneration. There is some evidence that a central role of PML nuclear bodies is the antiviral defence ^26,50^. Plenty of proteins are known to associate with PML bodies and nearly all are modified by SUMO and SUMOylation. The SUMO pathway is also relevant for the regulation of innate immune signalling during viral infection ^51^. Indeed, many viruses target PML bodies within infection. But what is the role of APP in this context? Notably, APP is capable to form homo-oligomers with the gp41 envelope protein from human immunodeficiency virus ^52^, co-purifies with Herpes simplex virus 1, and is also cotransported with HSV1 in living cells ^53,54^. The Kunitz protease inhibitor domain within the extracellular APP has sequence homology to Bikunin, which has been shown to interact with the Hepatitis E virus ORF3 ^55^. APP is found in all vertebrates and its molecular sequence shows a high degree of conservation, which is a typical feature of factors that significantly contribute to biological fitness. Thus, we speculate that APP acts as a viral response (receptor) protein that activates three spatially divided mechanisms involving the β- and γ-secretase driven cleavage producing β-amyloid and APP-CT (Figure 5). There is evidence from *in vitro* and *in vivo* studies for a role of ß-amyloid as antimicrobial peptide, e.g. it was shown that the peptide inhibits the replication of seasonal and pandemic strains of H3N2 and H1N1 influenza A virus (IAV) *in vitro* ^56^. Moreover, VZV-infection of primary human spinal astrocytes induce an extracellular amyloidogenic environment ^57^. Clinical studies reveal a deposition of amyloid-β plaques in the brains of HIV-infected individuals ^58^ and infections by HSV cause not only elevated β-amyloid production but also phosphorylation of Tau ^59,60^. Thus, β-amyloid might correspond to the first antiviral defence strategy in the extracellular matrix targeting the viral particle and lowering its infection potential (Figure 5). The second defence strategy, which is supported by our results, is the APP-dependent impact on the generation of PML bodies in the nucleus. This mechanism, including the fusion of APP-CT and PML aggregates to macro complexes, might modulate the cell for a viral nucleic acid invasion in the nucleus, thereby awakening the machinery to be prepared for the degradation of the pathogenic molecules in the APP-CT/PML aggregates. Reports showing an inclusion of DNA in PML bodies ^48^, the induction of PML by APP-CT ^19^ as well as findings of HSV-depending nuclear translocation of APP-CT ^61^ fit to this hypothesis. A reduction in nuclear PML bodies, as yet shown in high plaque areas of the human hippocampus, might restrict the cellular response potential to viral infection. Due to ongoing (repetitively over many years) infections, the nuclear defence strategy including the fusion of PML bodies is disturbed and as an alternative the cell needs to undergo programmed cell death in order to prevent virus propagation corresponding to a pyrrhic victory (Figure 5). Alternatively, the reduced PML bodies might correspond to an increased aggregation in infected cells starting with many tiny dots to a single macro-complex, and finally to nuclei without any PML signal, which potentially corresponds to dead cells. The third mechanism involves the phosphorylation of TAU, thereby reducing the intracellular transport of viral particles to replication sites. Notably, HSV-dependent TAU phosphorylation at multiple sites has been shown ^62^. There is strong evidence that reactivation of latent HSV is associated with dementia, but due to high latent infection in the population additional factors are needed to cause neuronal degeneration. A known AD susceptibility gene isoform is APOE □4, and notably, an association between HSV carriage and declined episodic memory function was described in individuals with APOE □4 genotype ^63^. Clinical correlation studies also revealed that for individuals with high frequency of HSV-1 reactivation (identified by immunoglobulin M (IgM) positiveness or elevated levels of IgG), the risk of developing AD is increased, whereas no significant association was found in APOE □4-negative subjects ^64^. A study on HIV-1 points to a potential mechanism of APOE in viral infection. Out of the three APOEs, APOE □4 was the least potent and effective at preventing HIV-1 internalization ^65^. To complete this model, it is highly likely that viral infection is accompanied by the activation of microglia cells causing inflammation including activation of TREM2 ^59,66,67^, corresponding to another AD susceptibility gene. Further research is necessary to underline this hypothetical model, which supports the benefit of an antiviral therapy for the treatment of AD as recently initiated ^68^.

**Figure 5.**
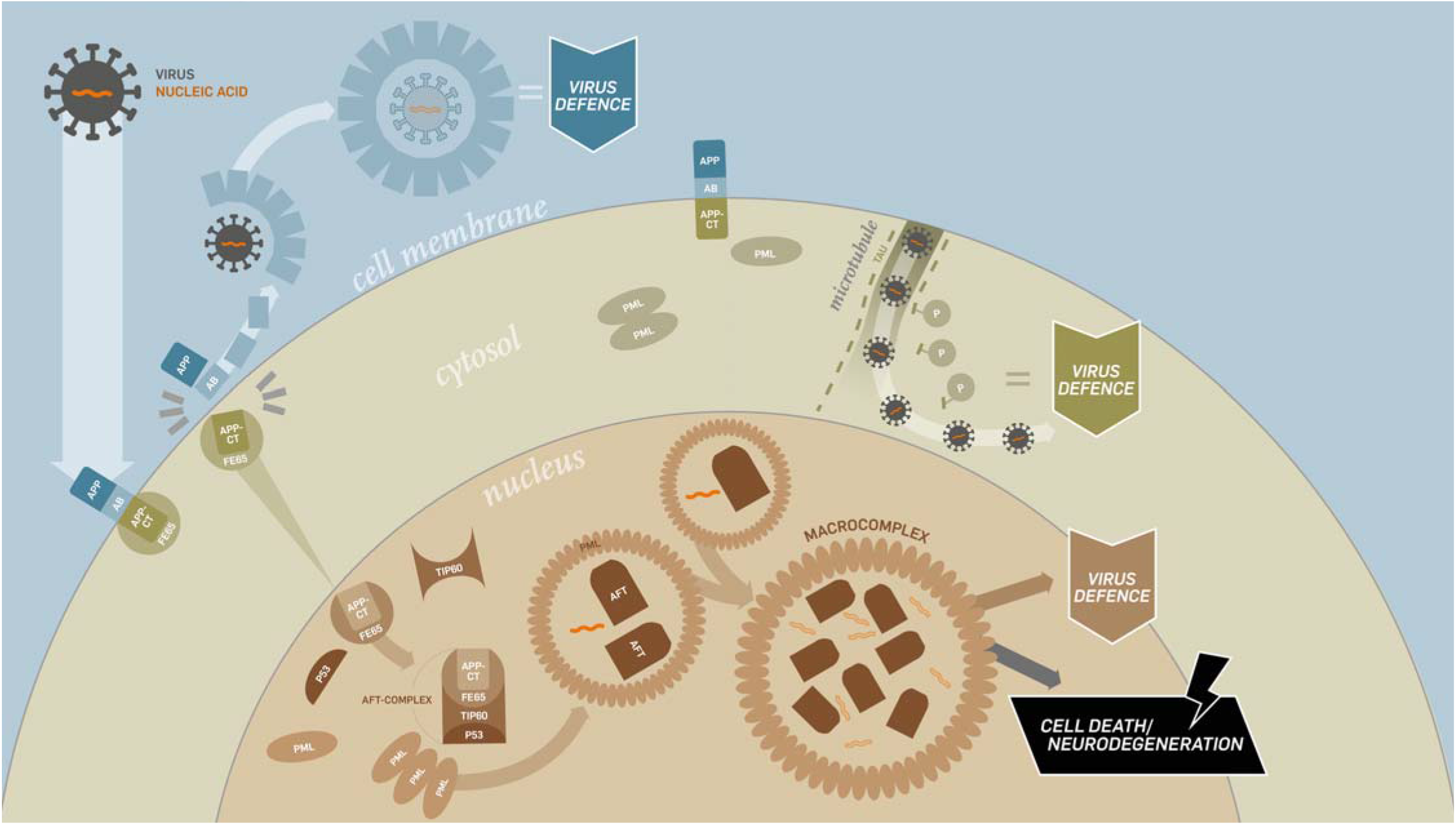
Hypothetical role of APP in virus defence. Our model suggests a twofold APP-related response to viral infection supported by our results and current literature as mentioned in the discussion. APP cleavage causes generation of β-amyloid, which has been shown to have antiviral relevance. The second defence mechanism is the sequestration of APP-CT, its signalling to the nucleus, and the generation of PML containing complexes. These AFT (APP-CT/FE65/TIP60)/PML aggregates might encapsulate viral nucleic acid preventing it from replication or initiating its degradation. However, overwhelming the response machinery due to repetitive infection (over years during live) or reduced immune system fitness might oblige the cell to degenerate as a last pyrrhic victory like defence strategy. Notably, the hyper-phosphorylation of TAU reducing the microtubule driven intracellular virus transport machinery might be associated to another virus defence strategy.

## Notes

### Competing Interest Statement

The authors have declared no competing interest.

## References

1. Hardy, J. & Selkoe, D.J. The amyloid hypothesis of Alzheimer’s disease: progress and problems on the road to therapeutics. Science 297, 353–356 (2002).

2. Bertram, L, Lili, C.M. & Tanzi, R.E. The genetics of Alzheimer disease: back to the future. Neuron 68, 270–281 (2010).

3. Lott, I.T., Head, E., Doran, E. & Busciglio, J. Beta-amyloid, oxidative stress and down syndrome. Curr Alzheimer Res 3, 521–528 (2006).

4. Chasseigneaux, S. & Allinquant, B. Functions of Aβ, sAPPα and sAPPβ: similarities and differences. J Neurochem 120 Suppl 1, 99–108 (2012).

5. Hardy, J.A. & Higgins, G.A. Alzheimer’s disease: the amyloid cascade hypothesis. Science 256, 184–185 (1992).

6. Bukhari, H., et al. Small things matter: Implications of APP intracellular domain AICD nuclear signaling in the progression and pathogenesis of Alzheimer’s disease. Prog Neurobiol 156, 189–213 (2017).

7. Karran, E. & De Strooper, B. The amyloid cascade hypothesis: are we poised for success or failure? J Neurochem 139 Suppl 2, 237–252 (2016).

8. Radzimanowski, J., et al. Structure of the intracellular domain of the amyloid precursor protein in complex with Fe65-PTB2. EMBO Rep 9, 1134–1140 (2008).

9. Schrötter, A., et al. FE65 regulates and interacts with the Bloom syndrome protein in dynamic nuclear spheres - potential relevance to Alzheimer’s disease. J Cell Sci 126, 2480-2492 (2013).

10. Grimm, M.O., et al. APP intracellular domain derived from amyloidogenic ß- and γsecretase cleavage regulates neprilysin expression. Front Aging Neurosci 7, 77 (2015).

11. Kim, H.S., et al. C-terminal fragments of amyloid precursor protein exert neurotoxicity by inducing glycogen synthase kinase-3beta expression. FASEB J 17, 1951–1953 (2003).

12. von Rotz, R.C., et al. The APP intracellular domain forms nuclear multiprotein complexes and regulates the transcription of its own precursor. J Cell Sci 117, 4435–4448 (2004).

13. Zhou, F., Gong, K., van Laar, T., Gong, Y. & Zhang, L. Wnt/β-catenin signal pathway stabilizes APP intracellular domain (AICD) and promotes its transcriptional activity. Biochem Biophys Res Commun 412, 68–73 (2011).

14. Bukhari, H., et al. Membrane tethering of APP c-terminal fragments is a prerequisite for T668 phosphorylation preventing nuclear sphere generation. Cell Signal 28, 1725–1734 (2016).

15. Domingues, S.C., et al. RanBP9 modulates AICD localization and transcriptional activity via direct interaction with Tip60. J Alzheimers Dis 42, 1415–1433 (2014).

16. Jowsey, P.A. & Blain, P.G. Fe65 Ser228 is phosphorylated by ATM/ATR and inhibits Fe65-APP-mediated gene transcription. Biochem J 465, 413–421 (2015).

17. Konietzko, U., et al. A fluorescent protein-readout for transcriptional activity reveals regulation of APP nuclear signaling by phosphorylation sites. Biol Chem (2019).

18. Langlands, H., Blain, P.G. & Jowsey, P.A. Fe65 Is Phosphorylated on Ser289 after UV-Induced DNA Damage. PLoS One 11, eO155O56 (2016).

19. Song, H., et al. Critical role of presenilin-dependent γsecretase activity in DNA damage-induced promyelocytic leukemia protein expression and apoptosis. Cell Death Differ 20, 639–648 (2013).

20. Waldron, E., et al. Increased AICD generation does not result in increased nuclear translocation or activation of target gene transcription. Exp Cell Res 314, 2419–2433 (2008).

21. Lallemand-Breitenbach, V. & de Thé, H. PML nuclear bodies: from architecture to function. Curr Opin Cell Biol 52, 154–161 (2018).

22. Van Damme, E., Laukens, K., Dang, T.H. & Van Ostade, X. A manually curated network of the PML nuclear body interactome reveals an important role for PML-NBs in SUMOylation dynamics. Int J Biol Sci 6, 51–67 (2010).

23. Dellaire, G. & Bazett-Jones, D.P. PML nuclear bodies: dynamic sensors of DNA damage and cellular stress. Bioessays 26, 963–977 (2004).

24. Chang, H.R., et al. The functional roles of PML nuclear bodies in genome maintenance. Mutat Res 809, 99–107 (2018).

25. Stadler, M., et al. Transcriptional induction of the PML growth suppressor gene by interferons is mediated through an ISRE and a GAS element. Oncogene 11, 2565–2573 (1995).

26. Scherer, M. & Stamminger, T. Emerging Role of PML Nuclear Bodies in Innate Immune Signaling. J Virol 90, 5850–5854 (2016).

27. Lancaster, M.A. & Knoblich, J.A. Generation of cerebral organoids from human pluripotent stem cells. Nat Protoc 9, 2329–2340 (2014).

28. Okada, Y. & Nakagawa, S. Super-resolution imaging of nuclear bodies by STED microscopy. Methods Mol Biol 1262, 21–35 (2015).

29. Cao, X. & Südhof, T.C. A transcriptionally [correction of transcriptively] active complex of APP with Fe65 and histone acetyltransferase Tip60. Science 293, 115–120 (2001).

30. Gersbacher, M.T., et al. Turnover of amyloid precursor protein family members determines their nuclear signaling capability. PLoS One 8, e69363 (2013).

31. Goodger, Z.V., et al. Nuclear signaling by the APP intracellular domain occurs predominantly through the amyloidogenic processing pathway. J Cell Sci 122, 3703–3714 (2009).

32. Hébert, S.S., et al. Regulated intramembrane proteolysis of amyloid precursor protein and regulation of expression of putative target genes. EMBO Rep 7, 739–745 (2006).

33. Edbauer, D., Willem, M., Lammich, S., Steiner, H. & Haass, C. Insulin-degrading enzyme rapidly removes the beta-amyloid precursor protein intracellular domain (AICD). J Biol Chem 277, 13389–13393 (2002).

34. Farris, W., et al. Insulin-degrading enzyme regulates the levels of insulin, amyloid beta-protein, and the beta-amyloid precursor protein intracellular domain in vivo. Proc Natl Acad Sci USA 100, 4162–4167 (2003).

35. Kimberly, W.T., Zheng, J.B., Guénette, S.Y. & Selkoe, D.J. The intracellular domain of the beta-amyloid precursor protein is stabilized by Fe65 and translocates to the nucleus in a notch-like manner. J Biol Chem 276, 40288–40292 (2001).

36. Walsh, D.M., et al. gamma-Secretase cleavage and binding to FE65 regulate the nuclear translocation of the intracellular C-terminal domain (ICD) of the APP family of proteins. Biochemistry 42, 6664–6673 (2003).

37. Fogal, V., et al. Regulation of p53 activity in nuclear bodies by a specific PML isoform. EMBO J 19, 6185–6195 (2000).

38. Wang, X.W., et al. Functional interaction of p53 and BLM DNA helicase in apoptosis. J Biol Chem 276, 32948–32955 (2001).

39. Tang, Y., Luo, J., Zhang, W. & Gu, W. Tip60-dependent acetylation of p53 modulates the decision between cell-cycle arrest and apoptosis. Mol Cell 24, 827–839 (2006).

40. Loe, T.K., et al. Telomere length heterogeneity in ALT cells is maintained by PML-dependent localization of the BTR complex to telomeres. Genes Dev (2020).

41. Duilio, A., et al. A rat brain mRNA encoding a transcriptional activator homologous to the DNA binding domain of retroviral integrases. Nucleic Acids Res 19, 5269–5274 (1991).

42. Kesavapany, S., et al. Expression of the Fe65 adapter protein in adult and developing mouse brain. Neuroscience 115, 951–960 (2002).

43. Simeone, A., et al. Expression of the neuron-specific FE65 gene marks the development of embryo ganglionic derivatives. Dev Neurosci 16, 53–60 (1994).

44. Hu, Q., Hearn, M.G., Jin, L.W., Bressler, S.L. & Martin, G.M. Alternatively spliced isoforms of FE65 serve as neuron-specific and non-neuronal markers. J Neurosci Res 58, 632–640 (1999).

45. Sun, Y., et al. A novel function of the Fe65 neuronal adaptor in estrogen receptor action in breast cancer cells. J Biol Chem 289, 12217–12231 (2014).

46. Squatrito, M., Gorrini, C. & Amati, B. Tip60 in DNA damage response and growth control: many tricks in one HAT. Trends Cell Biol 16, 433–442 (2006).

47. Vogt, D.L., et al. Abnormal neuronal networks and seizure susceptibility in mice overexpressing the APP intracellular domain. Neurobiol Aging 32, 1725–1729 (2011).

48. Luciani, J.J., et al. PML nuclear bodies are highly organised DNA-protein structures with a function in heterochromatin remodelling at the G2 phase. J Cell Sci 119, 2518–2531 (2006).

49. Meng, F., Na, I., Kurgan, L. & Uversky, V.N. Compartmentalization and Functionality of Nuclear Disorder: Intrinsic Disorder and Protein-Protein Interactions in Intra-Nuclear Compartments, hit J Mol Sci 17 (2015).

50. Tavalai, N. & Stamminger, T. New insights into the role of the subnuclear structure ND10 for viral infection. Biochim Biophys Acta 1783, 2207–2221 (2008).

51. Everett, R.D., Boutell, C. & Hale, B.G. Interplay between viruses and host sumoylation pathways. Nat Rev Microbiol 11, 400–411 (2013).

52. Abad, C., Martínez-Gil, L., Tamborero, S. & Mingarro, I. Membrane topology of gp41 and amyloid precursor protein: interfering transmembrane interactions as potential targets for HIV and Alzheimer treatment. Biochim Biophys Acta 1788, 2132–2141 (2009).

53. Cheng, S.B., Ferland, P., Webster, P. & Bearer, E.L. Herpes simplex virus dances with amyloid precursor protein while exiting the cell. PLoS One 6, e17966 (2011).

54. Satpute-Krishnan, P., DeGiorgis, J.A. & Bearer, E.L. Fast anterograde transport of herpes simplex virus: role for the amyloid precursor protein of alzheimer’s disease. Aging Cell 2, 305–318 (2003).

55. Tyagi, S., Surjit, M. & Lal, S.K. The 41-amino-acid C-terminal region of the hepatitis E virus ORF3 protein interacts with bikunin, a kunitz-type serine protease inhibitor. J Virol 79, 12081–12087 (2005).

56. White, M.R., et al. Alzheimer’s associated β-amyloid protein inhibits influenza A virus and modulates viral interactions with phagocytes. PLoS One 9, e101364 (2014).

57. Bubak, A.N., et al. Varicella-Zoster Virus Infection of Primary Human Spinal Astrocytes Produces Intracellular Amylin, Amyloid-β, and an Amyloidogenic Extracellular Environment. J Infect Dis 221, 1088–1097 (2020).

58. Hategan, A., et al. HIV Tat protein and amyloid-β peptide form multifibrillar structures that cause neurotoxicity. Nat Struct Mol Biol 24, 379–386 (2017).

59. Bortolotti, D., Gentili, V., Rotola, A., Caselli, E. & Rizzo, R. HHV-6A infection induces amyloidbeta expression and activation of microglial cells. Alzheimers Res Ther 11, 104 (2019).

60. Qin, Q. & Li, Y. Herpesviral infections and antimicrobial protection for Alzheimer’s disease: Implications for prevention and treatment. J Med Virol 91, 1368–1377 (2019).

61. Civitelli, L., et al. Herpes simplex virus type 1 infection in neurons leads to production and nuclear localization of APP intracellular domain (AICD): implications for Alzheimer’s disease pathogenesis. J Neurovirol 21, 480–490 (2015).

62. Wozniak, M.A., Frost, A.L. & Itzhaki, R.F. Alzheimer’s disease-specific tau phosphorylation is induced by herpes simplex virus type 1. J Alzheimers Dis 16, 341–350 (2009).

63. Lövheim, H., et al. Herpes Simplex Virus, APOE⍰4, and Cognitive Decline in Old Age: Results from the Betula Cohort Study. J Alzheimers Dis 67, 211–220 (2019).

64. Linard, M., et al. Interaction between APOE4 and herpes simplex virus type 1 in Alzheimer’s disease. Alzheimers Dement 16, 200–208 (2020).

65. Khan, N., Datta, G., Geiger, J.D. & Chen, X. Apolipoprotein E isoform dependently affects Tat-mediated HIV-1 LTR transactivation. J Neuroinflammation 15, 91 (2018).

66. Talwar, P., et al. Viral Induced Oxidative and Inflammatory Response in Alzheimer’s Disease Pathogenesis with Identification of Potential Drug Candidates: A Systematic Review using Systems Biology Approach. Curr Neuropharmacol 17, 352–365 (2019).

67. Carter, C.J. Alzheimer’s disease plaques and tangles: cemeteries of a pyrrhic victory of the immune defence network against herpes simplex infection at the expense of complement and inflammation-mediated neuronal destruction. Neurochem Int 58, 301–320 (2011).

68. Devanand, D.P., et al. Antiviral therapy: Valacyclovir Treatment of Alzheimer’s Disease (VALAD) Trial: protocol for a randomised, double-blind, placebo-controlled, treatment trial. BMJ Open 10, e032112 (2020).

