## Supplements for "Amyloid precursor protein causes fusion of promyelocytic leukemia nuclear bodies in human hippocampal areas with high plaque load"

Supplement

**Figure S1**

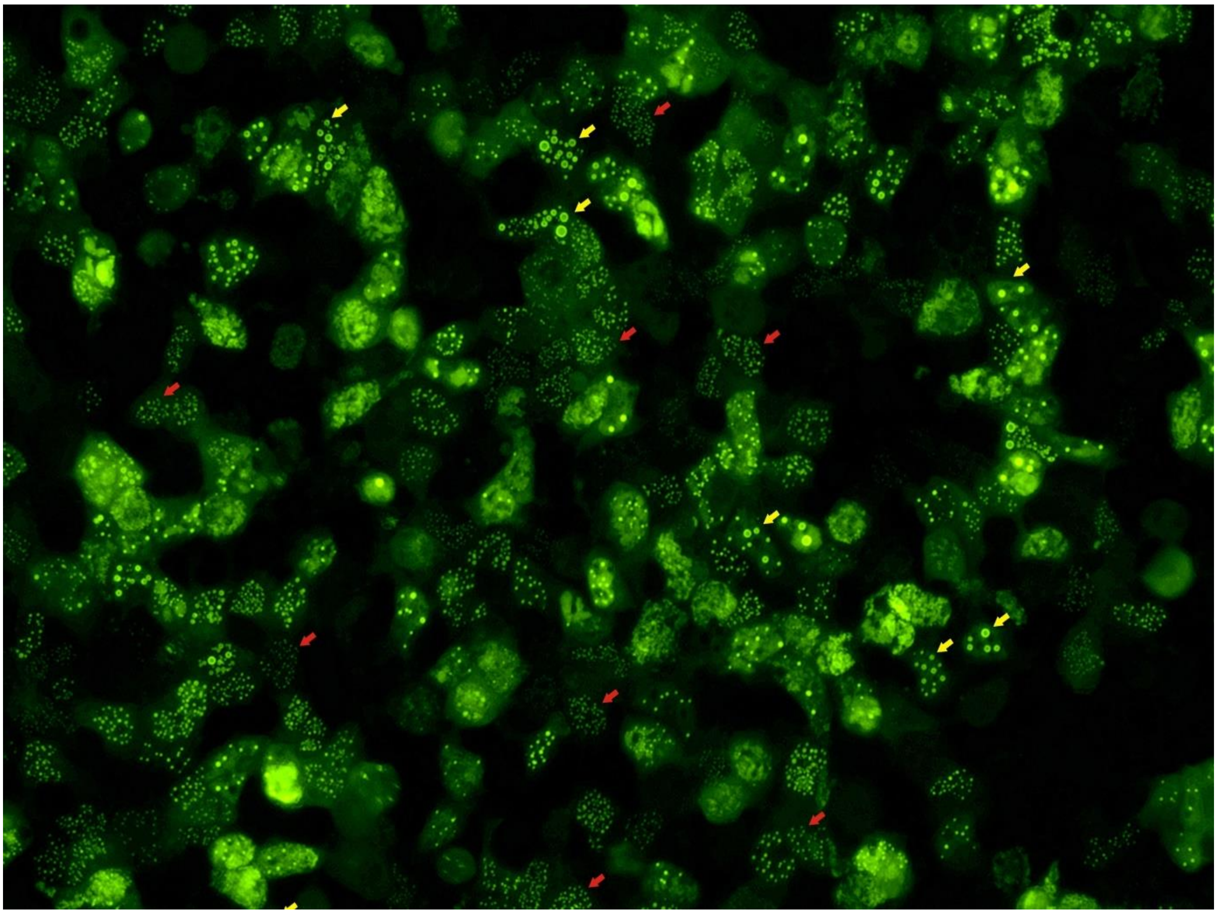

Figure S1 - HEK293 cells transfected with FE65-EGFP/TIP60 (w/o fluorophore) reveal a nuclear dot like phenotype. Cells with many tiny dots (red arrow) can be distinguished from cells with giant ring-like structures (yellow arrows).

**Figure S2**

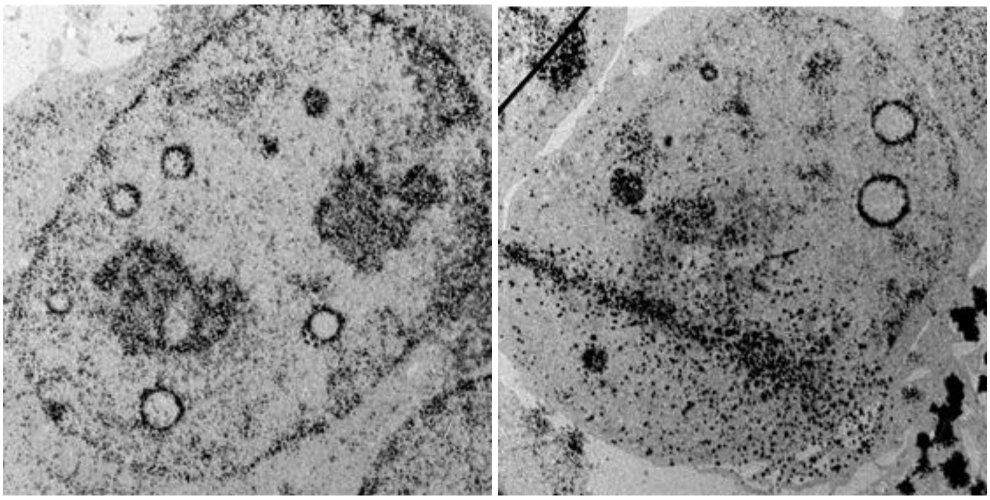

Figure S2 - Electron microscopy imaging of HEK293 cells, transfected with FE65-EGFP/TIP60 (w/o fluorophore) revealed a circular phenotype of the nuclear aggregates. However, membrane-like structured were not evident.

**Figure S3**

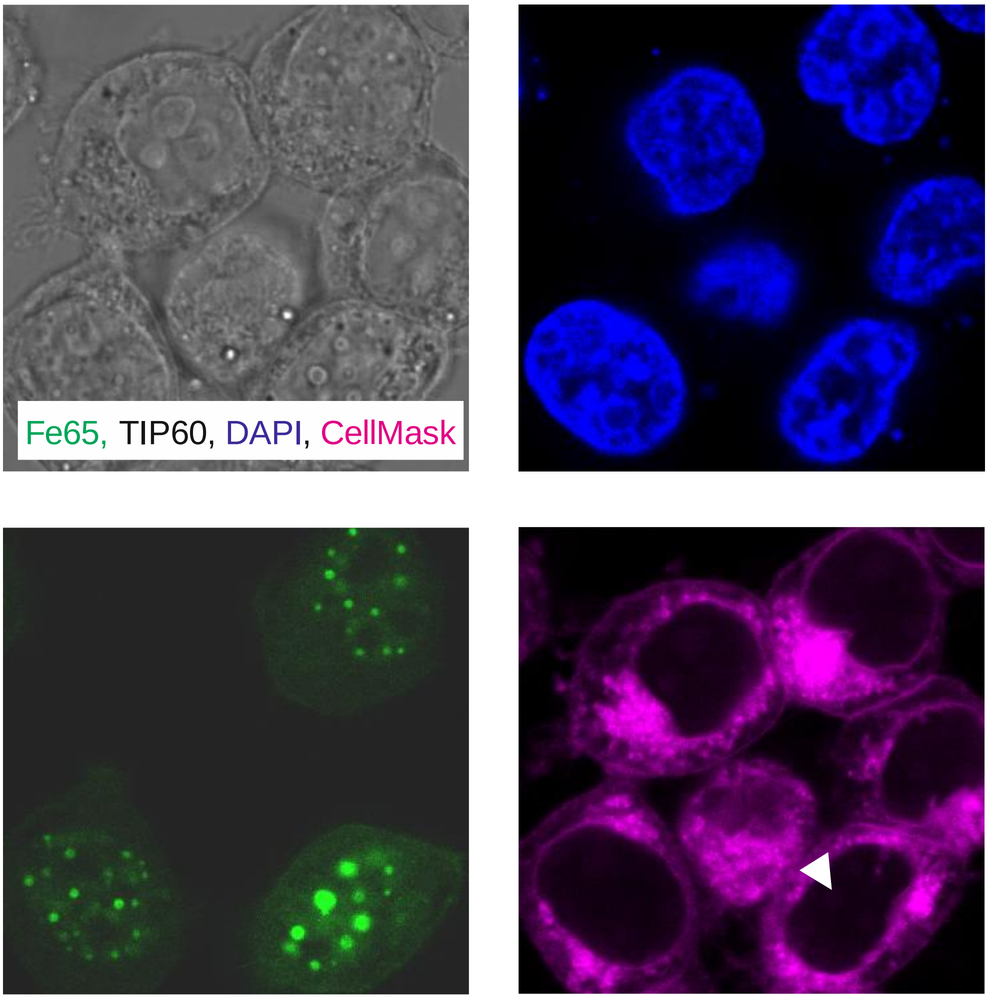


Figure S3 - CellMask stain (purple) in FE65-EGFP (green), TIP60 (w/o fluorophore) transfected HEK293 cells failed to reveal a prominent membrane staining encapsulating the nuclear aggregates (arrow).

**Figure S4**

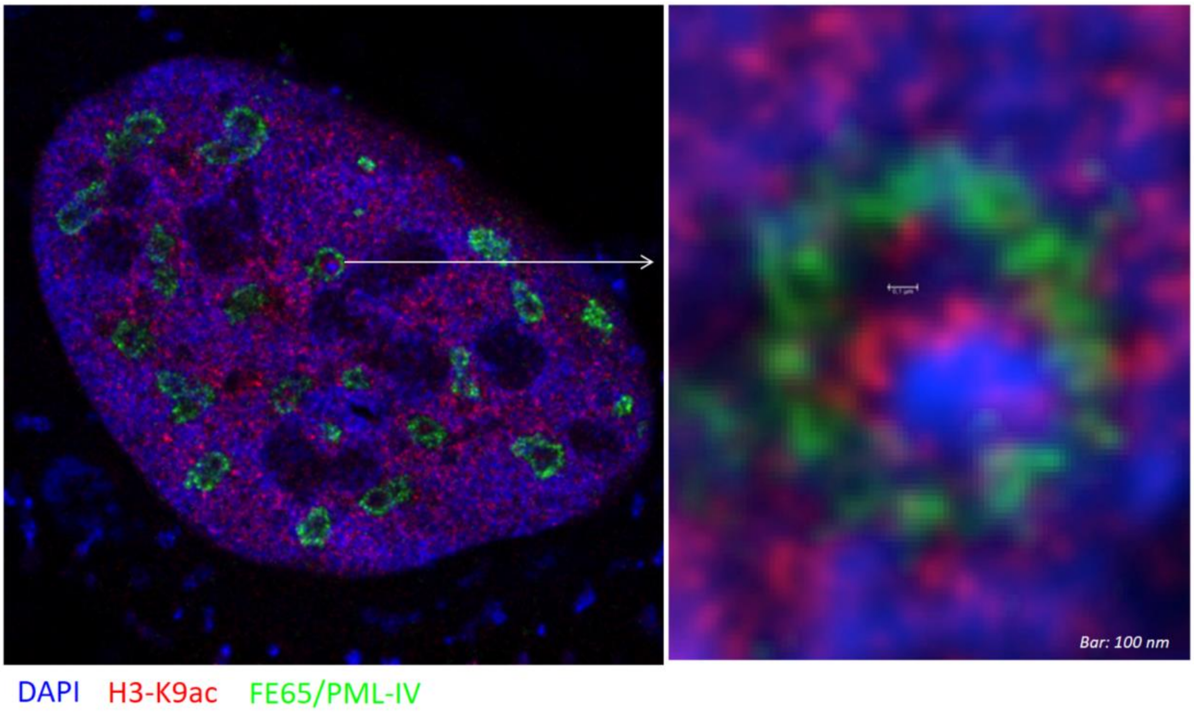


Figure S4 - STED imaging of FE65-EGFP, TIP60 (w/o fluorophore), PML (isoform IV) transfected cells. Upon fixation, immunofluorescence staining using anti histone3-K9ac antibody (red) was performed demonstrating active gene expression in some of the nuclear aggregates.

**Figure S5**

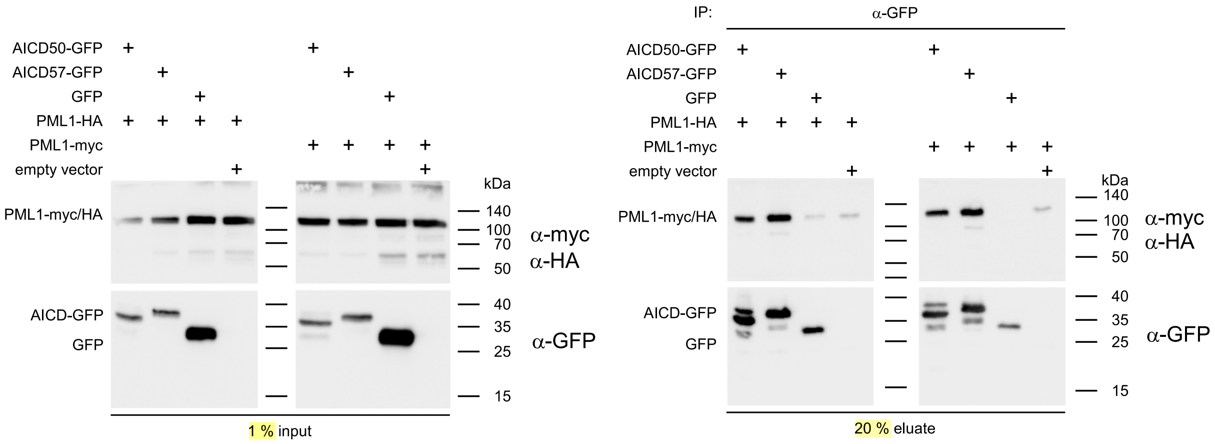


Figure S5 - Additional experiment to IP given in Figure 2K. For details reader is referred to Legend of Figure 2.

Figure S6

A

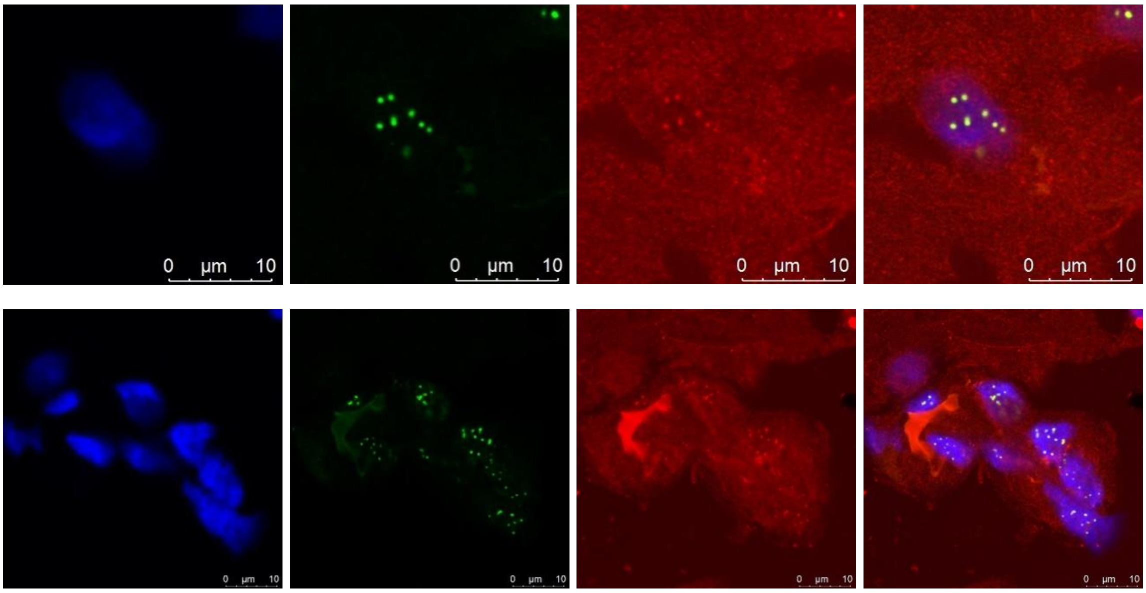


B

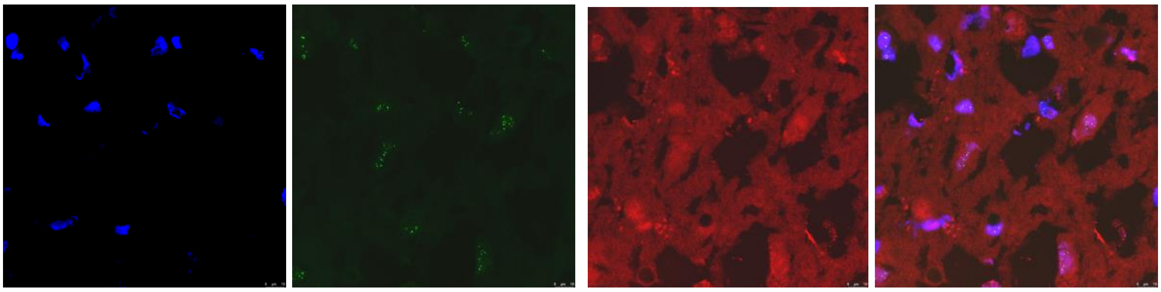

Figure S6 – APP-CT/PML co-staining in the human brain. (A) Additional images (to Figure 3E) revealing PML/APP-CT co-localization in human brain sections. (B) For negative control, the primary APP-CT antibody was omitted revealing no co-localization of the red (background) signal to PML in green.
